# scDREAMER: atlas-level integration of single-cell datasets using deep generative model paired with adversarial classifier

**DOI:** 10.1101/2022.07.12.499846

**Authors:** Ajita Shree, Musale Krushna Pavan, Hamim Zafar

**Author notes:** Authors contributed equally (Ajita Shree, Musale Krushna Pavan).

## Abstract

Integration of heterogeneous single-cell sequencing datasets generated across multiple tissue locations, time and conditions is essential for a comprehensive understanding of the cellular states and expression programs underlying complex biological systems. Here, we present scDREAMER (https://github.com/Zafar-Lab/scDREAMER), a novel data integration framework that employs a novel adversarial variational autoencoder for learning lower-dimensional cellular embeddings and a batch classifier neural network for the removal of batch effects. Using five real benchmarking datasets, we demonstrated that scDREAMER can overcome critical challenges including the presence of skewed cell types among batches, nested batch effects, large number of batches and conservation of development trajectory across different batches. Moreover, our benchmarking demonstrated that scDREAMER outperformed state-of-the-art methods in batch-correction and conservation of biological variation. Using a 1 million cells dataset, we further showed that scDREAMER is scalable and can perform atlas-level integration across species (e.g., human and mouse) while being faster than other deep-learning-based methods.

## Introduction

The exploration of cellular heterogeneity and developmental trajectories in different tissue systems has been revolutionized by the rapid advances of single-cell RNA-sequencing technologies (1; 2; 3). This rapid development coupled with large-scale collaborative initiatives such as the Human Cell Atlas (HCA) and Human BioMolecular Atlas Program (HuBMAP) (4; 5) have increased the complexity of single-cell datasets which can include samples contributed by different laboratories (6), generated across tissue locations, time and conditions (7; 8). Since single-cell datasets generated from similar biological contexts but different experimental conditions can share cellular features (9), integration of information from heterogeneous data sources can facilitate the discovery of major and rare cell types, improve the reconstruction of developmental trajectories (10) and lead to more reliable investigation of complex biological systems. However, the intrinsic differences in measured gene expression among experimental settings contributed by factors such as sequencing protocols, library preparation, sample donors, tissue of origin, sampling time and condition inevitably creates complex, nested batch effects which can diminish the value of data integration by confounding biological signals (11). Thus the development of computational data integration methods that can reliably eliminate the complex batch effects without undermining biological variations is a major challenge in scRNA-seq analysis (12).

Existing methods for integration of scRNA-seq datasets can be broadly classified into two groups. The first group of methods employ cell type annotations for supervised cross-batch learning (13; 14) while removing batch effects. The requirement for cell type annotations limits their applications as novel cell types cannot be captured. In comparison, unsupervised data integration methods (15; 16; 17; 18) that do not require cell type annotations are more widely used. Some of these methods identifies the batch-specific gene factors from the gene expression profiles (19). Methods such as BBKNN, Scanorama and Seurat v3 employ global mutual nearest neighbors (MNNs) i.e., paired cells between multiple batches for batch correction of the neighborhood using reduced-dimension cellular spaces. Harmony achieves better batch mixing by applying a novel local correction. These MNN-based methods consume a large memory and low-quality MNN can make it difficult to simultaneously identify dataset-specific cell types and the cell types that are shared by multiple datasets. Methods such as scVI (20) and DESC (21) that employ deep variational autoencoders for learning cellular embeddings from scRNA-seq can also integrate data from multiple batches. However, the traditional autoencoder models are challenged due to lower fidelity in reproducing the batch-corrected expression profiles (22). A recent benchmarking study (23) showed that methods such as Harmony and Seurat perform well for simple integration tasks but poorly for complex integration tasks and vice versa for Scanorama and scVI and a consistent tradeoff exists between batch-correction and preservation of biological variations.

To overcome the existing challenges, here we present a novel deep learning-based data integration framework, called scDREAMER (single cell Deep generative intRgrAtion Model with advErsarial classifieR) that employs a novel adversarial variational autoencoder consisting of an autoencoder and a discriminator and trained using evidence lower bound and Bhattacharyya loss. This network takes as input the high-dimensional scRNA-seq data and learns the lower-dimensional cellular embeddings that preserves the biological heterogeneity extricated from the technical variations. scDREAMER further incorporates a batch classifier (a multi-layer neural network) which is adversarially trained along with the encoder using a cross-entropy loss for the removal of batch effects.

We evaluated the performance of scDREAMER on 5 real datasets consisting of up to 1 million cells and 30 batches (Supplementary Table 1). For these complex data integration tasks, scDREAMER was able to overcome a variety of challenges such as the presence of skewed cell types among batches (pancreas integration), nested batch effects (lung integration), large number of batches (macaque retina integration) and scenarios when cell types of development trajectory are obtained from different batches (human immune integration). For all these integration tasks, scDREAMER achieved better performance in batch-correction and conservation of biological variation against that of the state-of-the-art methods. Using two different metrics, we also showed that scDREAMER can reliably identify rare cell types. scDREAMER further outperformed the other methods on another single-cell level metric that evaluates the efficacy of a method in discriminating batch-specific cell types and mixing shared cell types. Finally, using a 1 million cells dataset, we also demonstrated that scDREAMER is highly scalable, can perform atlas-level data integration across different species, and achieves runtime advantage over other deep-learning based integration methods.

## Results

### Overview of scDREAMER

Fig. 1 shows the overview of scDREAMER, which employs a novel adversarial variational autoencoder for learning the lower-dimensional representation of cells from the high-dimensional scRNA-seq data and a neural network classifier (also called a batch classifier) for removal of batch effects. scDREAMER models the scRNA-seq data as a nonlinear function of a lower-dimensional cell-state embedding and the batch information that encodes the variation in data generation. The adversarial variational autoencoder of scDREAMER consists of three multi-layer neural networks: an encoder *E* that maps the high-dimensional expression data (*x*_*i*_) and batch information (*s*_*i*_) of a cell *i* to a lower-dimensional embedding *z*_*i*_, a decoder *D*, which reconstructs the expression profile of the cell from *z*_*i*_ and *s*_*i*_, and a discriminator 𝒟 that aims to distinguish the original expression profile *x*_*i*_ and the expression profile reconstructed 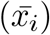 by the decoder. The adversarial variational autoencoder network of scDREAMER is trained using two loss functions: evidence lower bound is used for training the encoder and decoder networks, whereas Bhattacharyya loss is used for adversarial training of discriminator and autoencoder parameters. scDREAMER further incorporates a batch classifier ℬ (a multi-layer neural network) that takes as input the lower-dimensional embedding *z*_*i*_ learned by the encoder and tries to predict the batch information for cell *i*. The batch classifier and the encoder are adversarially trained using a cross-entropy loss where the encoder tries to maximize it with an aim to generate the embeddings such that the classifier is not able to differentiate between batches and the batch classifier tries to minimize it by distinguishing the embeddings of the cells coming from different batches and hence achieving better mixing of the batches (see Methods section for details).

**Figure 1:**
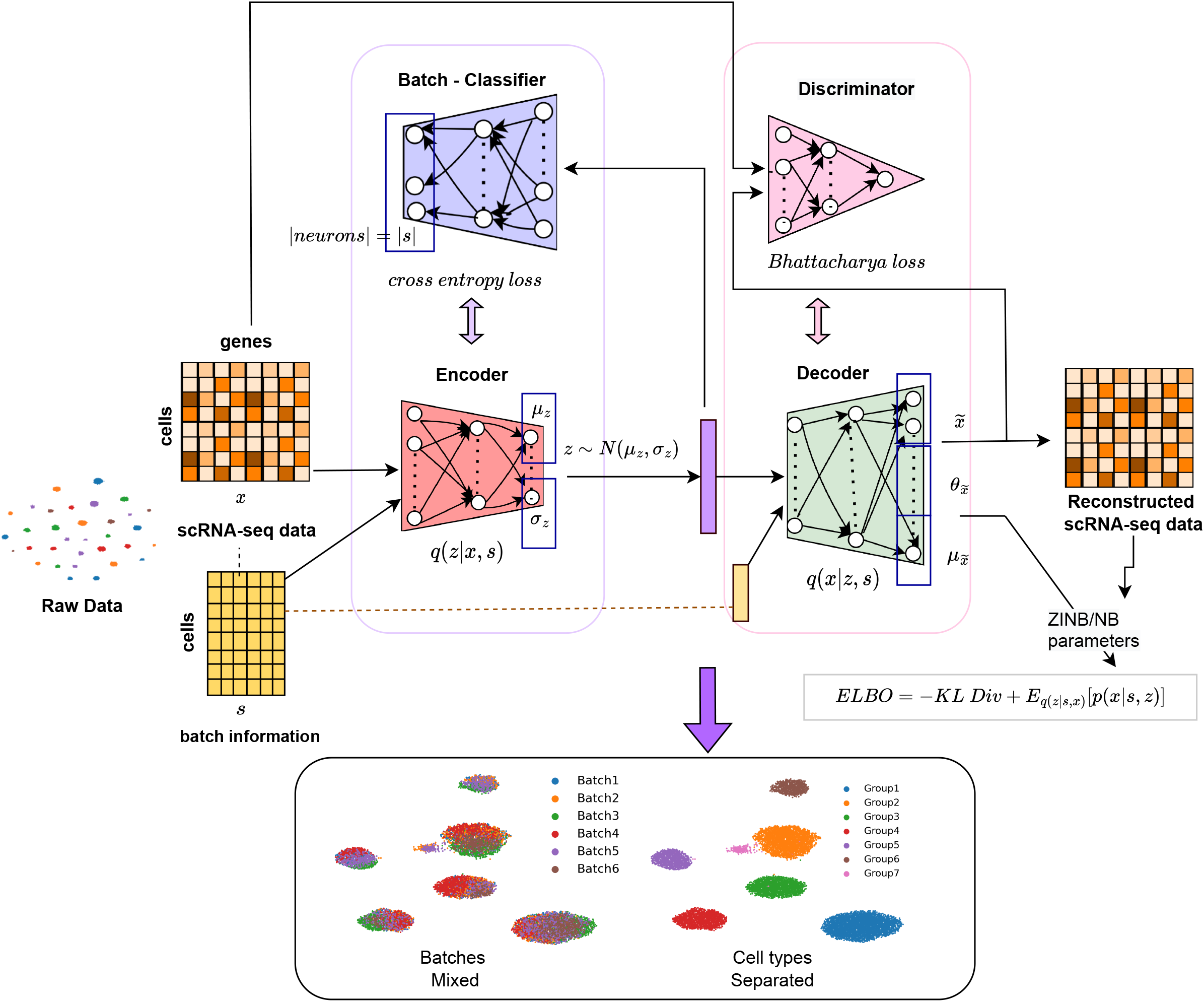
Overview of scDREAMER framework. scDREAMER consists of an adversarial variational autoencoder and a batch classifier. The adversarial variational autoencoder comprises of three networks: an encoder, a decoder, and a discriminator and these networks are trained using ELBO and Bhattacharya loss functions. The batch classifier is adversarially trained along with the encoder using a cross-entropy loss. scDREAMER learns latent cellular embeddings such that the cells from different batches are well-mixed and different cell types are separated leading to conservation of biological variations.

### Integration of pancreatic islet data generated using different sequencing protocols

We first tested scDREAMER’s ability to perform integration and batch correction across different sequencing protocols using a human pancreas dataset consisting of 16382 cells (Supplementary Fig. 1a). The dataset consisted of nine sub-datasets generated using distinct sequencing protocols including CEL-seq, CEL-seq2, Fluidigm C1, SMART-seq2, and inDrop. The dataset harbored 14 pancreatic cell types including acinar cells, activated and quiescent stellate cells, alpha cells, beta cells, delta cells, ductal cells, endothelial cells, epsilon cells, gamma cells, macrophages, mast cells, schwann cells and t cells. Integration was performed using scDREAMER and the performance was compared against that of six other integration methods. scDREAMER was able to almost perfectly separate all the cell types and mix the shared cell types across different protocols (Fig. 2a-b). In comparison, integration by scVI, BBKNN, Scanorama and INSCT led to restricted mixing of the different batches (Supplementary Fig. 1b-g). While Harmony and Seurat were also able to mix the batches well, Harmony was not able to clearly distinguish activated and quiescent stellate cells whereas Seurat improperly mixed some alpha cells with beta and ductal cells.

**Figure 2:**
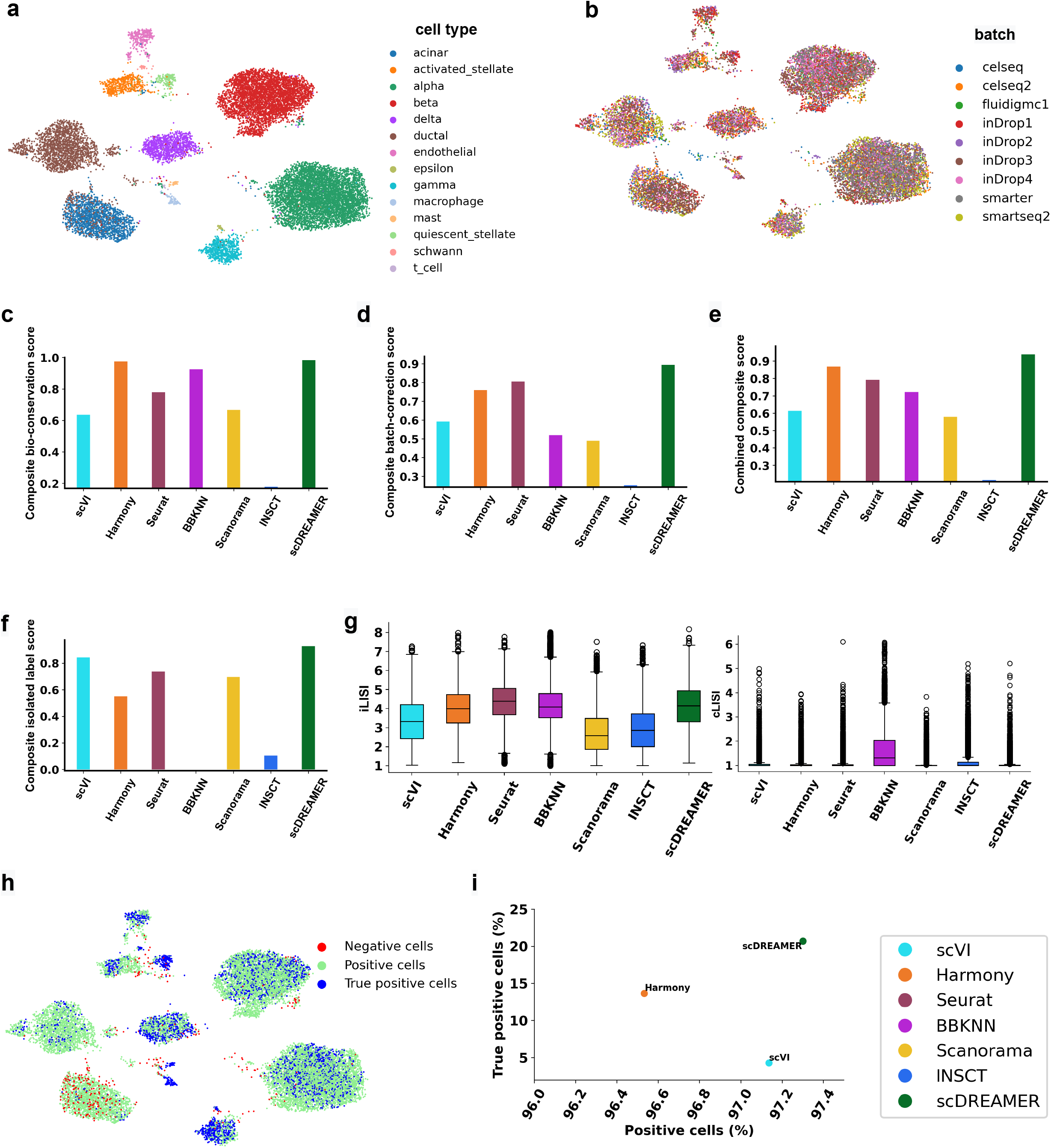
Integration of pancreatic islet data. (a) Visualization of scDREAMER’s latent space embeddings after integration of pancreatic islet dataset. Different colours denote different pancreatic cell types. (b) Visualization of scDREAMER’s latent space embeddings, cells are coloured based on batch information. Comparison of (c) composite bio-conservation score, (d) composite batch-correction score and (e) combined composite score metrics between scVI, Harmony, Seurat, BBKNN, Scanorama, INSCT and scDREAMER. (f) Comparison of composite isolated label scores to assess how well rare cell types are identified. (g) Comparison of iLISI and cLISI values. (h) Qualitative assessment of batch-mixing by visualization of scDREAMER’s latent space embeddings, cells are coloured based on three categories - positive, negative and true positive. (i) Quantitative assessment of batch-mixing of scDREAMER against scVI and Harmony based on the percentage of positive vs true positive cells.

Next, we quantitatively compared scDREAMER’s performance against that of the other methods based on four composite accuracy scores. Composite bio-conservation score measures the accuracy of a method in preserving biological variance after integration and considers global clustering accuracy (normalized mutual information (NMI) and Adjusted Rand Index (ARI)) and relative distances between clusters (cell type average silhouette width (ASW)). The accuracy of batch effect removal was measured using composite batch-correction score which considers four different metrics including the k-nearest-neighbor batch effect test (kBET), ASW across batches, k-nearest-neighbor (kNN) graph connectivity and batch removal using PCA regression. Combined composite score computes the average of composite bio-conservation and composite batch-correction score. We also used a composite isolated label score that evaluates the ability of a method in capturing rare cell identities based on f1 score and silhouette coefficient in identifying the rare cells.

scDREAMER consistently outperformed all other methods by achieving the highest combined composite score which was driven by scDREAMER’s superior performance in both conservation of biological variance and batch correction (Fig. 2c-e). While scVI, Harmony, and Scanorama performed similar to scDREAMER in terms of NMI and ARI metrics, scDREAMER and Harmony performed much better in terms of cell type ASW (Supplementary Fig. 2a). scDREAMER achieved the best composite batch-correction score because of its superior performance in terms of multiple batch-correction metrics (Supplementary Fig. 2b). scDREAMER was also the best method in capturing the rare cell identities as it achieved the best isolated label scores (Fig. 2f, Supplementary Fig. 2c). While Harmony was the second best method in terms of combined composite score, it performed poorly in terms of composite isolated label score indicating its inability to capture the rare cell identities correctly. We also measured the local inverse Simpson’s Index scores (iLISI for batch mixing and cLISI for cell-type separation) at single-cell level and based on these metrics, scDREAMER’s performance was comparable to that of Harmony and Seurat and better than all other methods (Fig. 2g). We further adopted another single-cell level evaluation measure which computes the proportion of “positive” (cells connected to other cells only from the same cell type) and “true positive” (fraction of positive cells whose local and global batch distributions are congruous) cells after integration and used these summary metrics to compare scDREAMER’s performance against that of Harmony (second best method in terms of combined composite score) and scVI (second best method in terms of composite isolated label score). Again, scDREAMER outperformed both Harmony and scVI based on the proportion of positive and true positive cells (Fig. 2h-i).

### Integration of lung atlas consisting of cells from different human donors

Next, we applied scDREAMER for the integration of lung atlas data consisting of 32472 cells from lung transplant and biopsy samples from 16 donors sequenced using 10X and drop-seq (Supplementary Fig. 3a). The integration of this dataset poses several challenges including inter-donor variability, protocol-specific batch effects (10X samples A1-A6, 1-6; drop-seq samples B1-B4), and variability across sampling type and tissue locations. Specifically, cell type composition varied between the transplant samples (1-6, B1-B4) that are obtained from lung parenchyma and the biopsy samples (A1-A6) obtained from lung airways. Two basal cell types, ciliated and secretory cells were majorly present in the biopsy samples but the transplant samples harbored them only as minor populations or in some cases the cell types were absent. The dataset also contained rare cell types present across a few donors (ionocytes). Furthermore, the endothelial and secretory cells should vary across biopsy and transplant samples due to transcriptome being affected by tissue location.

scDREAMER was able to successfully integrate the batches across sequencing protocols overcoming donor-level variation (Fig. 3a-b, Supplementary Fig. 4a-b). Only scDREAMER and scVI were clearly able to identify rare ionocytes as a separate cluster, whereas the other methods mixed them with other cell types. scDREAMER was able to identify all biopsy-specific cell types and also preserved basal cell subtypes. The secretory cells from the biopsy and transplant samples were separated by scDREAMER (Supplementary Fig. 4a). The lymphatic and endothelial cells were incorrectly merged by Harmony, Seurat and INSCT whereas Scanorama, scVI and scDREAMER was able to identify them as separate clusters (Supplementary Fig. 3b-g). The endothelial cells were separated into three clusters by Scanorama, two of which corresponded to transplant samples indicating incomplete integration of these samples. scDREAMER was able to preserve the spatial variation of endothelial cells at higher resolution. Neutrophil subpopulations were merged by all the methods.

**Figure 3:**
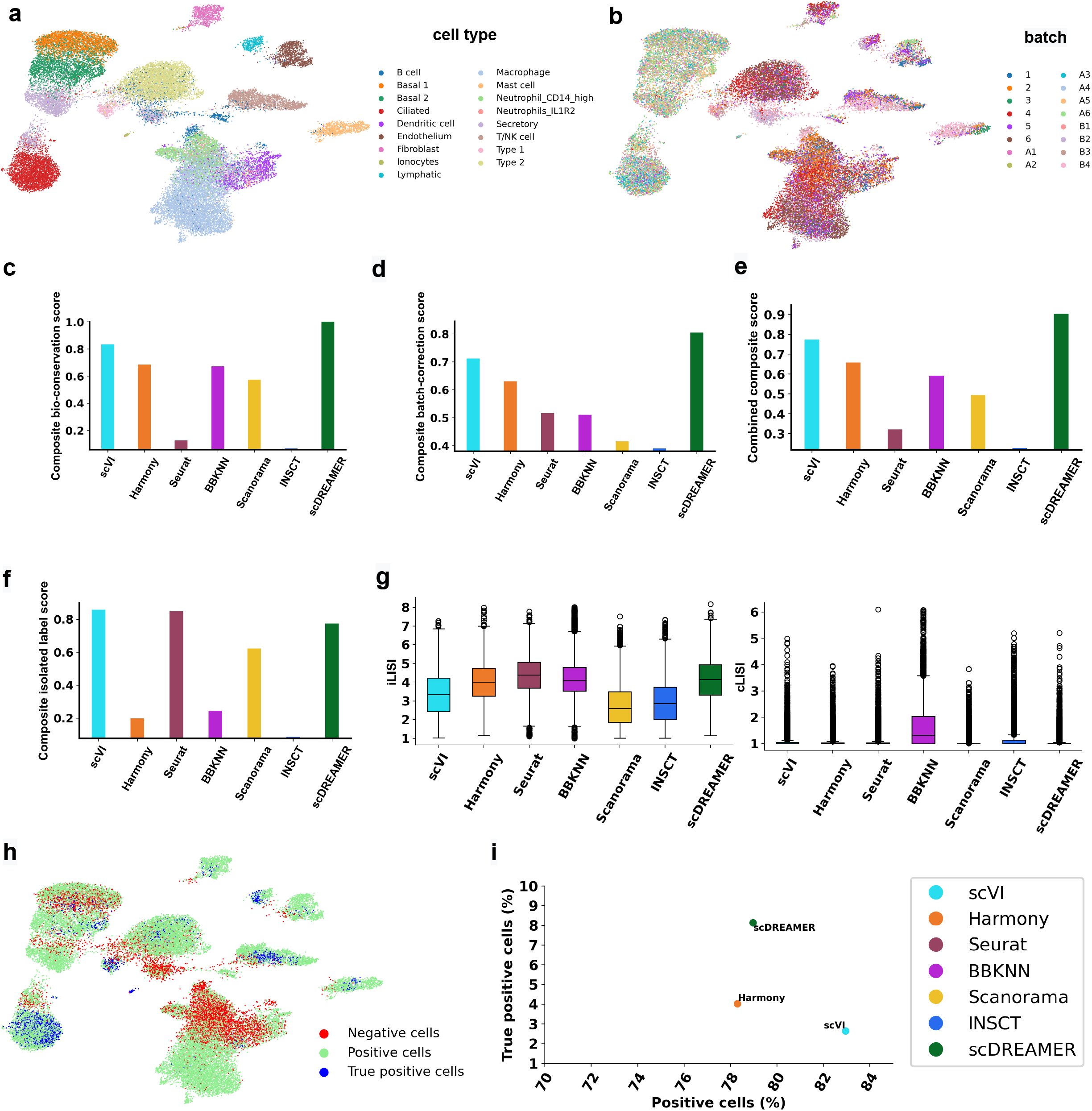
Integration of lung atlas data. (a) Visualization of scDREAMER’s latent space embeddings after integration of lung atlas dataset. Different colours denote different lung cell types. (b) Visualization of scDREAMER’s latent space embeddings, cells are coloured based on the batch information. Comparison of (c) composite bio-conservation score, (d) composite batch-correction score and (e) combined composite score metrics between scVI, Harmony, Seurat, BBKNN, Scanorama, INSCT and scDREAMER for the integration of lung atlas data. (f) Comparison of composite isolated label scores to assess how well rare cell types are identified. (g) Comparison of iLISI and cLISI values. (h) Qualitative assessment of batch-mixing by visualization of scDREAMER’s latent space embeddings, cells are coloured based on three categories - positive, negative and true positive. (i) Quantitative assessment of batch-mixing of scDREAMER against scVI and Harmony based on the percentage of positive vs true positive cells.

scDREAMER performed superior to all other methods both in terms of bio-conservation and batch correction achieving the highest combined composite score (Fig. 3c-e). scDREAMER out-performed all the methods in terms of all three bio-conservation metrics (Supplementary Fig. 5a). While scVI performed similar to scDREAMER in terms of PCR Batch, graph connectivity and ASW label/batch metrics, scDREAMER and Harmony performed much better in terms of kBET (Supplementary Fig. 5b). In capturing rare cell identities, scDREAMER performed comparably with the top performing methods (Fig. 3f, Supplementary Fig 5c), in fact it achieved the highest isolated f1 score. While scVI was the second best method in terms of combined composite score, its performance was poor for batch mixing. While Harmony was the third best method in terms of combined composite score, it performed poorly in terms of composite isolated label score indicating its inability to capture the rare cell identities correctly. In terms of LISI metrics, scDREAMER performed the best in terms of cLISI and comparably to scVI and Harmony in terms of iLISI metrics outperforming all other metrics (Fig. 3g). We further compared scDREAMER’s performance against that of scVI and Harmony (second and third-best methods based on combined composite score respectively) by measuring the proportion of positive and true positive cells. Due to the close spacing of dendritic cells and Neutrophil subtypes in the embedding, we observed the presence of large number of negative cells for these cell types (same observation for the embedding of other methods). While scVI had the highest proportion of positive cells, scDREAMER outperformed both the methods based on the proportion of true positive cells (Fig. 3h-i) without sacrificing much in terms of positive proportion.

### Integration of human immune cells from peripheral blood and bone marrow of different donors

We next evaluated scDREAMER’s integration and batch correction performance for integrating 33506 human immune cells obtained in ten batches corresponding to different donors and the cells were sampled from bone marrow and peripheral blood (PBMCs) and the sequencing was performed using two protocols (10X and smart-seq2) (Supplementary Fig. 6a). Out of 33506 cells, 9581 cells were sampled from bone marrow and comprised of three batches (Oetjen et al. (24)), and the rest 23985 bone marrow cells comprised of seven batches (10X genomics (25), Freytag (26), Sun et al (27) batches and Villani et al. (28)). While the cells in (28) were sequenced using smart-seq2, all other batches were sequenced using 10X genomics protocol. This integration task poses several challenges including donor-level variability, sequencing protocol-specific batch effects and cell types spanning multiple tissues. In addition, the dataset also contained cell subtypes that are difficult to distinguish because of transcriptional similarity (e.g., CD8+ and CD4+ T cells; CD14+ and CD16+ monocytes) and some tissue-specific cell types (e.g., monocyte progenitors, erythroid progenitors, erythrocytes and CD10+ B cells were only present in bone marrow) that need to be identified as separate clusters. Finally, the dataset also harbored the developmental trajectory of erythrocytes from hematopoietic stem and progenitor cells (HSPCs) via megakaryocyte and erythroid progenitors and the conservation of this trajectory across batches is an important aspect of this integration task.

scDREAMER was able to resolve the inter-sample and inter-platform batch effects while integrating human immune dataset as indicated by disjoint well-mixed cell type clusters (Fig. 4a-b). scDREAMER was successful in clustering together the same cell types across tissues while identifying cell subtypes, such as CD8+ and CD4+ T cells, CD20+ and CD10+ B cells, CD14+ and CD16+ monocytes, as distinct closely located clusters. In comparison, CD20+ and CD10+ B cells were distantly placed in BBKNN and Scanorama embeddings (Supplementary Fig 6f-g). While scDREAMER was able to place monocyte-derived dendritic cells and plasmacytoid dendritic cells in separate clusters in close vicinity (Fig. 4a-b), other methods such as Harmony, Seurat and INSCT (Supplementary Fig. 6c-d, 6g) placed these cell sub types far apart in the embedding. Furthermore, scVI separated monocyte-derived dendritic cells into multiple clusters, one of which was mixed with other distinct cell types (e.g., megakaryocyte progenitors, plasma cells, CD4+ T cells, etc.) indicating poor clustering (Supplementary Fig. 6b). Finally, scDREAMER clearly separated tissue-specific cell types and exhibited a perfect continuum of bone marrow cell types after integration, preserving the trajectory from HSPCs to erythrocytes (Fig. 4a-b, Supplementary Fig. 7a-b). In contrast, methods (Supplementary Fig. 6d, 6g) such as Seurat and INSCT failed to conserve the trajectory of erythrocyte development.

**Figure 4:**
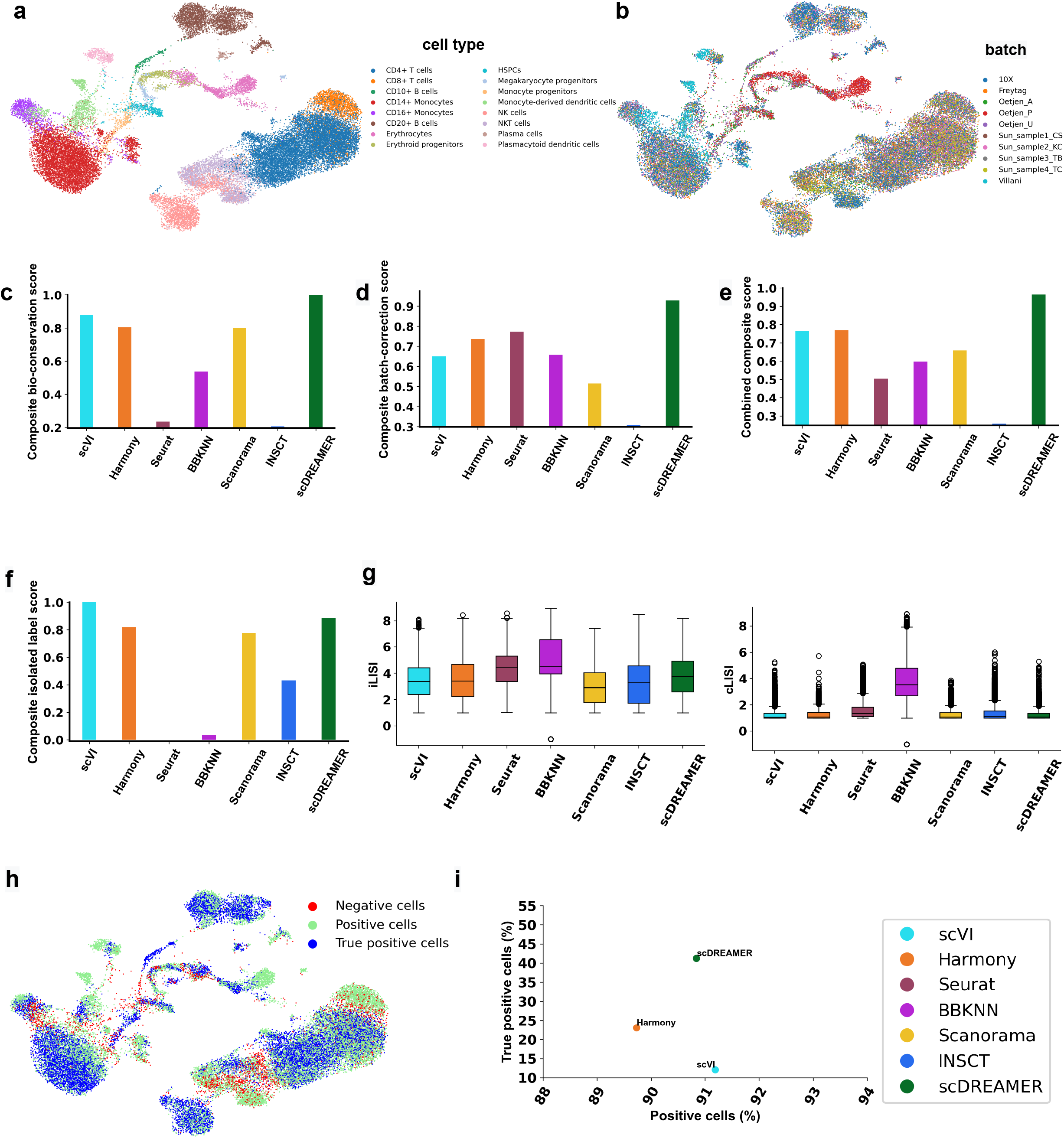
Integration of human immune data. (a) Visualization of scDREAMER’s latent space embeddings after integration of human immune dataset. Different colours denote different cell types present in the human immune dataset. (b) Visualization of scDREAMER’s latent space embeddings, cells are coloured based on the batch information. Comparison of (c) composite bio-conservation score, (d) composite batch-correction score and (e) combined composite score metrics between scVI, Harmony, Seurat, BBKNN, Scanorama, INSCT and scDREAMER for the integration of human immune data. (f) Comparison of composite isolated label scores to assess how well rare cell types are identified. (g) Comparison of iLISI and cLISI values. (h) Qualitative assessment of batch-mixing by visualization of scDREAMER’s latent space embeddings, cells are coloured based on three categories - positive, negative and true positive. (i) Quantitative assessment of batch-mixing of scDREAMER against scVI and Harmony based on the percentage of positive vs true positive cells.

Our quantitative comparison showed that scDREAMER consistently outperformed all other methods by achieving the highest combined composite score which was driven by scDREAMER’s superior performance in both biological conservation and batch correction (Fig. 4c-e). scDREAMER was the best performing method in terms of PCR batch, kBET and all three bio-conservation metrics (Supplementary Fig. 8a-b). In capturing the rare cell identities, scDREAMER also performed at par with scVI, the best method in terms of composite isolated label scores (Fig. 4f, Supplementary Fig. 8c). However, for batch correction scVI performed poorly. In terms of LISI metrics, scDREAMER along with scVI and Harmony consistently performed better than other methods (Fig. 4g). We further compared scDREAMER’s performance against that of Harmony and scVI (second and third-best methods based on combined composite score respectively) by measuring the proportion of positive and true positive cells. While both scDREAMER and scVI had comparable proportion of positive cells better than Harmony, scDREAMER outperformed both the methods by a large margin based on the proportion of true positive cells (Fig. 4h-i).

### Integration of macaque retina bipolar cells across a large number of batches

Next, we tested scDREAMER’s ability to integrate cells across a large number of batches by analyzing a dataset that contained 30302 bipolar cells from macaque retina (29) (Supplementary Fig. 9a). This dataset harbored 12 different cell types that originated from fovea or periphery of retina and consisted of 30 batches across macaques and regions. The original study showed that several integration methods failed to remove batch effects from this dataset (29).

scDREAMER successfully integrated all 30 batches while separating the distinct cell types (Fig. 5a-b). While scVI, Harmony, INSCT and BBKNN (Supplementary Fig. 9b-g) also captured the cell types in distinct cluster, scVI incorrectly placed some DB2 cells near rod bipolar cells, whereas Harmony, INSCT and BBKNN could not properly distinguish OFFx bipolar cells. In comparison, cell types were splitted into multiple clusters by Scanorama (Supplementary Fig. 9f).

**Figure 5:**
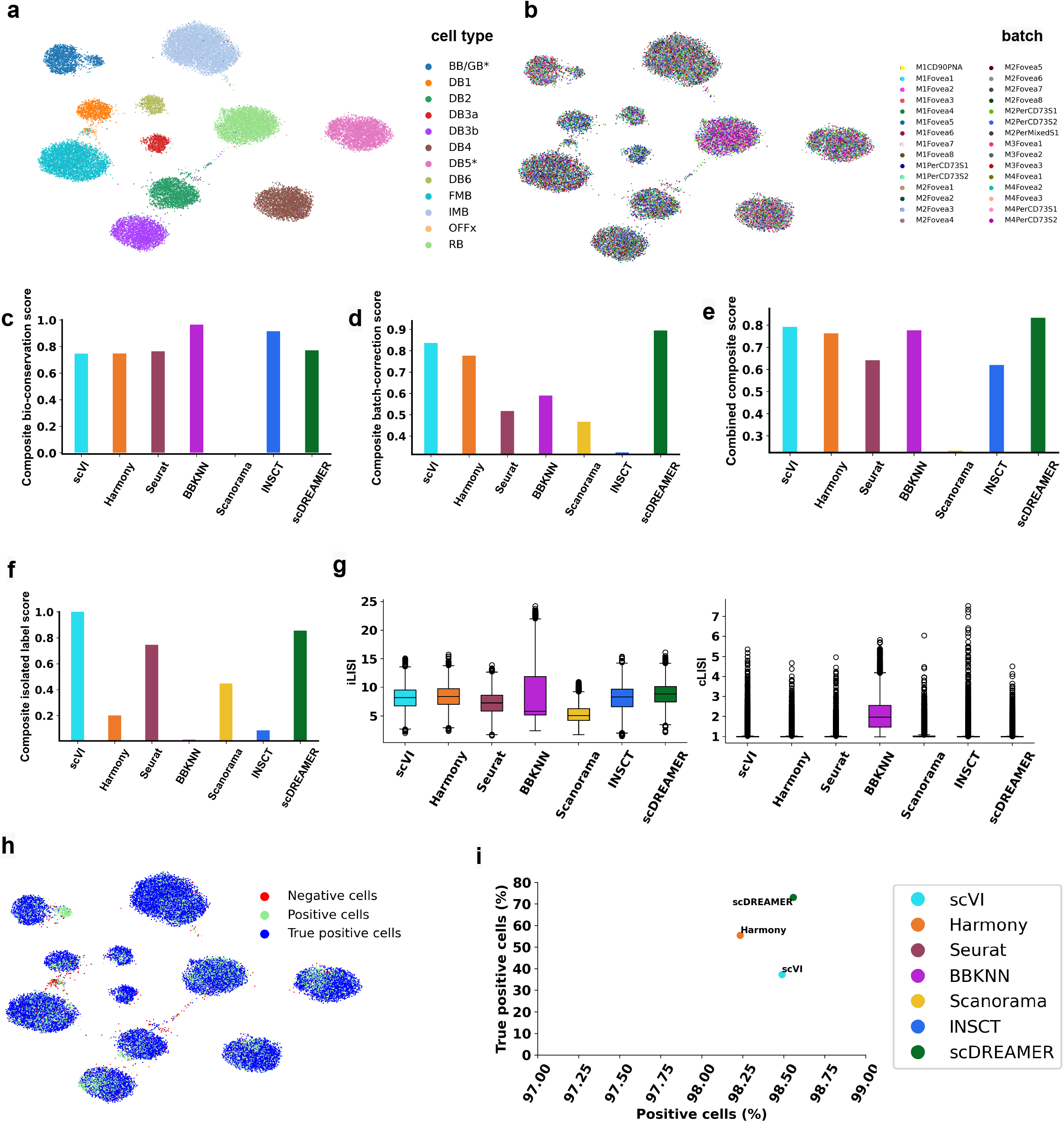
Integration of macaque retina bipolar cells. (a) Visualization of scDREAMER’s latent space embeddings after the integration of macaque retina bipolar cells. Different colours denote different retina biploar cell types. (b) Visualization of scDREAMER’s latent space embeddings, cells are coloured based on the batch information. Comparison of (c) composite bio-conservation score, (d) composite batch-correction score and (e) combined composite score metrics between scVI, Harmony, Seurat, BBKNN, Scanorama, INSCT and scDREAMER for the integration of macaque retina data. (f) Comparison of composite isolated label scores to assess how well rare cell types are identified. (g) Comparison of iLISI and cLISI values. (h) Qualitative assessment of batch-mixing by visualization of scDREAMER’s latent space embeddings, cells are coloured based on three categories - positive, negative and true positive. (i) Quantitative assessment of batch-mixing of scDREAMER against scVI and Harmony based on the percentage of positive vs true positive cells.

Quantitative analysis showed that scDREAMER was the top performer based on the combined composite score (8% improvement over the second-best method) which was driven by scDREAMER’s superior performance in batch correction (Fig. 5c-e). For biological conservation, scDREAMER achieved the best NMI and ARI values (Supplementary Fig. 10a). scDREAMER was the best performing method in terms of PCR batch and kBET batch correction metrics (Supplementary Fig. 10b). In capturing the rare cell identities, scDREAMER was the second-best performing method after scVI (Fig. 5f, Supplementary Fig. 10c). In terms of LISI metrics, scDREAMER and Harmony performed the best followed by scVI and INSCT (Fig. 5g). We further compared scDREAMER’s performance against that of Harmony and scVI by measuring the proportion of positive and true positive cells. scDREAMER had the highest proportion of positive cells. While scVI had a comparable proportion of positive cells to that of scDREAMER, it was outperformed by scDREAMER by a large margin based on the proportion of true positive cells (Fig. 5h-i).

### Robust integration of millions of cells across species using scDREAMER

We finally evaluated scDREAMER’s ability to perform atlas-level integration across species using a dataset consisting of ~ millions of cells from human and mouse profiled by the Human Cell Landscape (HCL) (30) and Mouse Cell Atlas (MCA) (31) projects respectively. The two batches in the dataset correspond to HCL and MCA respectively. The dataset harbored 97 different cell types which had minimal overlap across the atlases (Supplementary Fig. 11a). The HCL batch comprised of 599926 cells from 63 different cell types whereas mouse cell atlas comprised of 333778 cells from 52 different cell types. Total 18 cell types (among the 97 cell types) were common between these two atlases. We have demonstrated scDREAMER’s scalability over million cells and versatility in integration with this atlas-level integration task.

For this atlas-level integration task, the performance of scDREAMER was compared with that of scVI, Scanorama, Harmony, INSCT and BBKNN. Due to the size of the dataset, Seurat ran into memory issue. Qualitatively, we can observe that clusters by scDREAMER were better separated compared to other methods (Fig. 6a-b, Supplementary Fig. 11b-g). scDREAMER was able to clearly distinguish some of the major cell types such as neutrophil cells, erythroid cells, fetal stromal cells and oligodendrocyte cells (Fig. 6a). While scVI could cluster some of these cell types such as neutrophil, erythroid and fetal stromal cells well, harmony could only cluster erythroid cells. In Harmony embeddings, similar cell types were disjointly present in multiple clusters. On the other hand, BBKNN led to complete mixing of multiple cell types. While Scanorama was able to better classify than BBKNN and Harmony, it was not able to properly mix the batches as shown in (Supplementary Fig. 11e).

**Figure 6:**
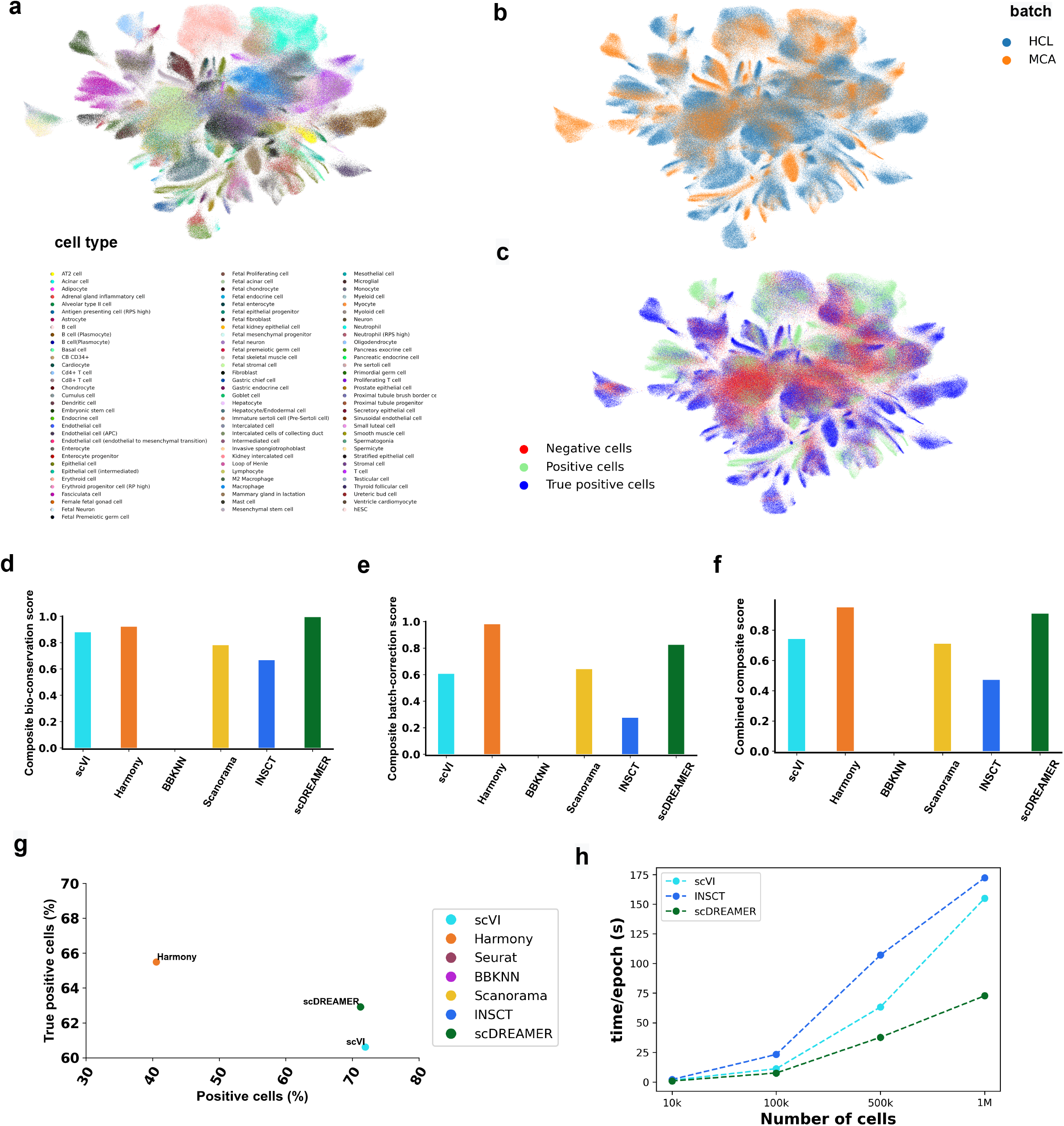
scDREAMER enables robust integration of millions of cells across species. (a) Visualization of scDREAMER’s latent space embeddings after the integration of human (HCL) and mouse cells (MCA). Different colours denote different cell types in this large dataset consisting of ~ million cells. (b) Visualization of scDREAMER’s latent space embeddings, cells are coloured based on the batch information. (c) Qualitative assessment of batch-mixing by visualization of scDREAMER’s latent space embeddings, cells are coloured based on three categories - positive, negative and true positive. Comparison of (d) composite bio-conservation score, (e) composite batch-correction score and (f) combined composite score metrics between scVI, Harmony, Seurat, BBKNN, Scanorama, INSCT and scDREAMER for the integration of HCL and MCA cells. (g) Quantitative assessment of batch-mixing of scDREAMER against scVI and Harmony based on the percentage of positive vs true positive cells. (h) Comparison of scDREAMER runtime against that of scVI and INSCT across four different scRNA datasets consisting of 10k, 100k, 500k, and 1M cells subsampled from the cross-species integration dataset.

Next, we quantitatively compared scDREAMER’s performance against that of the other methods based on three composite accuracy scores (Fig. 6d-f). scDREAMER and Harmony achieved the highest combined composite score. scDREAMER performed the best in terms of biological conservation and second-best in terms of batch correction (Fig. 6d-e, Supplementary Fig. 12a-b). For all three biological conservation metrics, scDREAMER performed the best (Supplementary Fig. 12a). We further compared scDREAMER’s performance against that of Harmony and scVI by measuring the proportion of positive and true positive cells. Based on the proportion of positive cells, both scDREAMER and scVI (> 71%) outperformed Harmony (40.5%) by a large margin. scDREAMER further outperformed scVI based on the proportion of true positive cells (Fig. 6c, 6g).

To assess scDREAMER’s scalability with the number of cells, we measured the runtime of scDREAMER on different datasets sub-sampled from the atlas-integration dataset and compared the runtime against that of scVI and INSCT - two other neural network-based methods. For all subsampled datasets, scDREAMER outperformed the other two methods based on runtime and exhibited higher scalability with the number of cells (Fig. 6h). For the complete dataset consisting of ~ 1 million cells, training scDREAMER took only ~ 72 seconds per epoch as compared to ~ 154 seconds per epoch for scVI and ~ 172 seconds per epoch for INSCT (on a server with one Nvidia Quadro RTX 5000 GPU).

## Discussion

Here, we introduced scDREAMER, a novel deep generative model for the efficient and robust integration of scRNA-seq datasets across multiple batches. scDREAMER employs a novel adversarial variational autoencoder for inferring the latent cellular embeddings from the high-dimensional gene expression matrices from different batches. This adversarial autoencoder also outputs the corrected expression profiles. The other component of scDREAMER, a batch classifier helps remove batch effects from the latent cellular embeddings for better mixing of cell types shared across multiple batches.

Our comprehensive benchmarking of scDREAMER on multiple complex data integration tasks demonstrate scDREAMER’s superiority over the state-of-the-art methods in terms of both conservation of biological variations and removal of batch effects. In contrast, the other methods were only able to perform well in one aspect of data integration: batch-mixing or conservation of biological variation, that too varied across datasets. scDREAMER was also a consistent performer in capturing the rare cell identities. Moreover, comparison of the fraction of true positive and positive cells further demonstrated scDREAMER’s superiority over other methods in batch-mixing.

scDREAMER also demonstrated high accuracy for the integration of atlases from different species despite the small number of shared cell types. Moreover, the unsupervised deep learning approach of scDREAMER does not require any cell type information and can be applied when prior knowledge regarding homologous cell types is not available or the cell type annotations are missing. As more cell atlases are generated from different species, we believe that scDREAMER will be suitable for robust integration of cross-species datasets for the discovery of shared and private cell types.

The application of scDREAMER on the cross-species dataset also highlights its scalability to millions of cells. In fact, our runtime experiments using different downsampled versions of this dataset showed that scDREAMER outperformed other deep learning-based methods based on runtime. Thus scDREAMER provides the most suitable deep learning-based integration model for cross-species atlas integration as the other deep learners scVI and INSCT achieved much less accuracy for cross-species integration task. This is particularly important as the deep learning-based methods enable the inference of latent cellular embeddings as well as corrected expression profiles which are required for several downstream applications such as trajectory inference (10) or differential expression analysis (20).

scDREAMER being a deep learning-based method may require fine-tuning of some parameters for extracting the best performance. However, in our analyses, same parameter values performed well for multiple datasets (Supplementary Table 2). Our current model assumes prior knowledge on the number of batches. An important future direction would be to explore the unsupervised treatment of batch information and whether the hierarchical structure between different batch information can be utilized when exists (e.g., when cells from a single donor but multiple organs are present in the atlas). While we restricted our analysis for the integration of scRNA-seq datasets, our deep generative model encompasses a general framework which can accommodate other omics datasets and we plan to extend the framework of scDREAMER to multiomics datasets. Finally, given the rapid generation of atlas-level single-cell datasets (32; 33; 34) across multiple organs and species, we anticipate scDREAMER to become an invaluable method for performing scalable and accurate integration of single-cell atlases for the exploration of different biological systems.

## Methods

### The scDREAMER model

scDREAMER consists of an unsupervised deep learning-based framework specifically designed to address the complex and multi-level batch effects and perform atlas-level integration ensuring effective integration as well as batch mixing. Any integration method is faced with the critical challenge of balancing a tradeoff between capturing the distinct identity of the batch-specific cell types and adequate mixing of the cell types shared across multiple batches. This challenge is even more profound for the integration of atlas-level datasets. To overcome these challenges, we formalize our scDREAMER integration model into two major components, the adversarial variational autoencoder and the batch classifier that is adversarially trained with the encoder.

scDREAMER employs a novel adversarial variational autoencoder for learning a lower-dimensional representation of cells from the high-dimensional scRNA-seq data. In addition, there is a neural network classifier (also called a batch classifier) for the removal of batch effects. The adversarial variational autoencoder of scDREAMER consists of three multi-layer neural networks: an encoder *E* that maps the high-dimensional expression data (*x*_*i*_) and batch information (*s*_*i*_) of a cell *i* to a lower-dimensional embedding *z*_*i*_, a decoder *D*, which reconstructs the expression profile of the cell from *z*_*i*_ and *s*_*i*_, and a discriminator 𝒟 that aims to distinguish the original expression profile *x*_*i*_ and the expression profile reconstructed 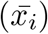 by the decoder. The use of a discriminator to adversarially train the autoencoder is inspired from (35).

The adversarial variational autoencoder network of scDREAMER is trained using two loss functions: evidence lower bound (ELBO) is used for training the encoder and decoder networks, whereas Bhattacharyya loss is used for adversarial training of discriminator and autoencoder parameters. The ELBO loss accounts for the KL divergence between the posterior distribution *q*(*z*_*i*_|*s*_*i*_, *x*_*i*_) and the true distribution *p*(*z*_*i*_), and the expected likelihood of *x*_*i*_ given *z*_*i*_ over posterior probability distribution *q*(*z*_*i*_|*s*_*i*_, *x*_*i*_). The Bhattacharyya loss compares the probability distributions *q*(*z*_*i*_|*x*_*i*_, *s*_*i*_) and *p*(*z*_*i*_) where *q*(*z*_*i*_|*x*_*i*_, *s*_*i*_) is the posterior probability distribution and *p*(*z*_*i*_) is 0-mean Gaussian distribution i.e. 𝒩(0, *I*).

scDREAMER further incorporates a batch classifier, ℬ (a multi-layer neural network) that takes as input the lower-dimensional embedding *z*_*i*_ learned by the encoder and tries to predict the batch information *s*_*i*_ for each cell *i*. The batch classifier and the encoder are adversarially trained using a cross-entropy loss where the encoder tries to maximize it with an aim to generate the embeddings such that the classifier is not able to differentiate between batches and the batch classifier tries to minimize it by distinguishing the embeddings of the cells that are part of different batches and hence achieving better mixing of the batches.

### Adversarial variational autoencoder for representation learning of cells

*E* denotes the Encoder network that takes as input *X* (gene expression matrix) and *S* (set of batch information) and generates the mean *µ*_*Z*_ and variance *σ*_*Z*_ of the multivariate normal distribution (prior for *z*). In addition it also learns the mean and variance of the cell-specific scaling factor *l*_*i*_ as used in (20). The following functional form is learnt by the Encoder.

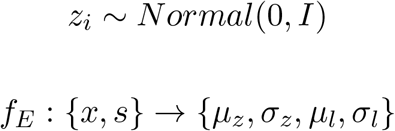

The decoder network *D* takes as input the latent space embeddings *z* and batch information and reconstructs the gene expression matrix, 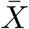. It also outputs the mean and dispersion of the reconstructed gene expression vector. The scRNA-seq data can be modeled as zero-inflated negative binomial (ZINB) or negative binomial (NB) distribution.

scDREAMER aims to learn the posterior distribution of the latent variable *z, p*(*z*|*X, S, l*). We use variational inference approach where we try to learn the network parameters of *E* and *D* as denoted by *ϕ* = {*f*_*E*_, *f*_*D*_} by maximizing the Evidence Lower Bound Loss (ELBO) function:

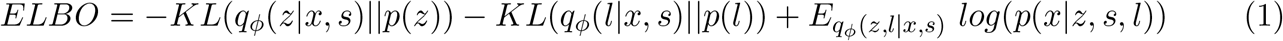

where, the first two terms denote the KL divergence between the posterior distribution and true distribution for *z* and *l* respectively and the third term denotes the expected likelihood of *x* given *z* over the posterior probability distribution, *q*_*ϕ*_(*z*|*s, x, l*). In this work we have modeled scRNA-seq data using ZINB distribution which is defined in terms of mean (*µ*) and dispersion (*θ*) parameters as:

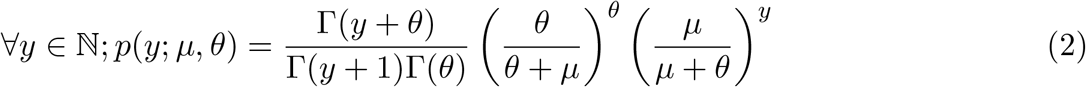

### Training of discriminator

Inspired by (35), scDREAMER autoencoder also incorporates an adversarial discriminator 𝒟 that tries to distinguish between the reconstructed gene expression profile 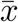 and the original gene expression profile *x* of a cell. This ensures that the distribution of the reconstructed gene expression profiles faithfully follow the distribution of the underlying original scRNA-seq profiles. 𝒟 is adversarially trained along with the autoencoder, 𝒜_ℰ_ = {*E, D*} using Bhattacharyya loss, ℒ_ℬ_ given by:

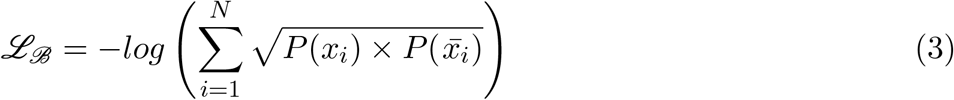

𝒟 tries to maximize ℒ_ℬ_ whereas the autoencoder 𝒜_ℰ_ tries to minimize ℒ_ℬ_ giving rise to the minimax objective function:

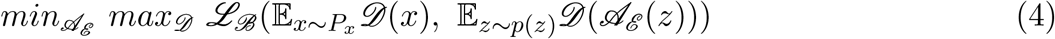

### Adversarial training using a batch classifier for batch-effect removal

The batch-classifier network ℬ learns its parameters by minimizing the cross entropy loss thereby ensuring correct classification of embeddings into different batches while Encoder trying to fool the batch-classifier by maximizing the cross entropy loss. ℬ takes *z* (latent space embeddings) as input and *s* (batch information) as labels. The cross entropy loss is given by:

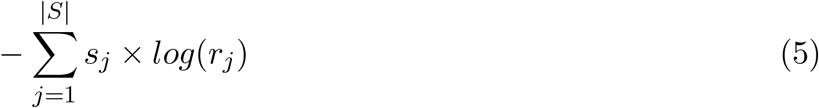

where *s*_*i*_ is the true label and *r*_*j*_ is the softmax probability of the *j*^*th*^ batch.

The batch-classifier intends to minimize the cross-entropy loss, hence classifying the latent cellular embeddings to its correct batch whereas the encoder tries to maximize the cross-entropy loss ensuring that the classifier is not able to differentiate between batches so that better mixing of data from different batches can be achieved. This gives rise to another minimax objective:

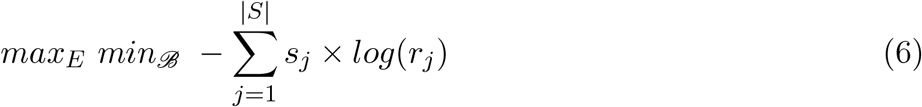

### Implementation details

- scDREAMER has 4 neural network units correspnding to Encoder, Decoder, Discriminator and Batch-Classifier, all with multi-layer dense neural network architecture with linear relu activation at the end of each layer. For training our model, we have adopted ADAM optimizer (36). We have used *β* as a scaling factor (multiplied to the KL-divergence term) to balance between the reconstruction loss and KL-divergence term while optimizing the ELBO loss. The batch size of 128 is used while training the model.
- To facilitate and stabilize the training process and make scDREAMER robust to small perturbations, we added a penalty term in the main objective function following (37). Specifically, for each gene expression vector *x*_*i*_, we down-sample *x*_*i*_ by keeping 80% of its UMIs to produce *x*_*i*_. The latent representations for *x*_*i*_ and 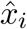 are *z*_*i*_ and 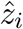 respectively. The penalty term is defined as 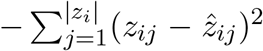 as we want *z*_*i*_ and 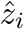 to be as close as possible. The down sampling ratio used in our case is 80%.

### Preprocessing of scRNA-seq datasets

The scRNA-seq datasets were preprocessed using the standard pipeline of Scanpy (38). Raw count expression matrices were imported as Scanpy AnnData object followed by the removal of low-quality cells based on the mitochondrial gene counts. The expression matrices were then normalized (using “scanpy.pp.normalize total” function) and log-transformed. Finally, we selected top 2000 highly variable genes using the function “scanpy.pp.highly variable genes” (with flavour parameter as “seurat”) as input to scDREAMER and other methods. The genes that were present across all the batches were considered.

### Metrics for the evaluation of integration performance

The data integration performance of all methods were primarily evaluated based on two broad categories of metrics: (1) biological variance conservation metrics and (2) batch effect correction metrics. Following a recent study (23), we considered NMI, ARI and ASW (cell type) as biological conservation metrics. For the evaluation of batch effect removal, we considered four metrics: ASW (batch), principal component regression (batch), graph connectivity, and kBET. For a holistic comparison of the performance across all the metrics in both of these categories, we introduced two composite scores and a combined composite score as described below. In addition, we computed isolated label F1 and isolated label silhouette scores for evaluating a method’s ability to capture rare cell identities. We also computed single-cell level metrics: graph iLISI (for batch correction), graph cLISI (for bio-conservation), and proportion of true positive vs positive cells.

### Biological conservation metrics

The computation of the biological conservation metrics requires clustering of the integrated data. We have used Louvain clustering algorithm to compute the clusters with a resolution that maximizes the NMI value. The same clustering assignment is used for computing other metrics (23).

- **Normalized Mutual Information (NMI):** NMI compares the overlap between any two clustering assignments where the overlap is measured in terms of mutual information. The NMI value is scaled in the range 0 to 1 based on the mean entropy for cluster assignments and cell-type labels.
- **Adjusted Rand Index (ARI):** Rand index compares two clustering assignments considering all pairs of points and finding agreement and disagreement between the clustering assignments (39). The adjusted rand index (ARI) accounts for the randomly correct labels and it ranges from 0 to 1 with 0 corresponding to random clustering, and 1 representing a perfect match between the clusterings (40). For ARI calculation, we compared the cell labels with the optimized NMI Louvain clusters.
- **Cell type average silhouette width (ASW):** Silhouette width for cell *i* is given by:

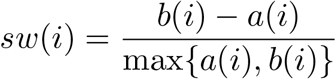

where *a*(*i*) is the average of distances of cell *i* to the other cells in the same cluster and *b*(*i*) is the average distance of cell *i* from the cells in the other nearest cluster. Average silhouette width (ASW) is calculated by averaging over all cells and it ranges between –1 to 1, where *−*1 or 0 indicates that clusters are overlapping and 1 corresponds to well separated and dense clusters. As a bio-conservation metric, we computed cell type ASW by considering the cell types as the clusters and scaled between 0 to 1 by the transformation ASW = (ASW + 1)/2. The PCA reduced space is used for the calculation of distances for the corrected space output method. This metric is not applicable to graph-based method BBKNN.

### Batch correction metrics

- **Batch ASW:** ASW is calculated between the batches of each subset based on the cell type. Here the 0 value indicates that the batches are well mixed. So we scaled the *ASW* = 1–*abs*(*ASW*). Now the scaled value 0 indicates that the batches are not well mixed, 1 indicates that batches are well mixed.
- **Principal component regression (PCR) batch:** In principal component regression (PCR) batch (41), for each principal component (*PC*_*k*_) (calculated using PCA), the variance contributed by each batch *b* is computed as:

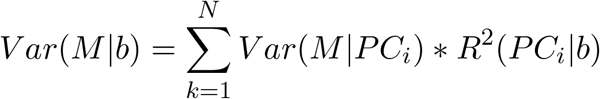 Where *V ar*(*M*|*PC*_*i*_) indicates the variance explained by *PC*_*i*_ on the data matrix *M*, and *R*^2^ denotes coefficient of determination calculated using a linear regression with *PC*_*i*_ as the dependent and *b* as the independent variable.
- **Graph Connectivity** Graph connectivity measures whether cells from the same cell type are well connected in the kNN graph. First, we compute the kNN graph with all the cells. Then we create a subset kNN graph with the cells from a particular cell type and check the number of cells in its largest connected component.

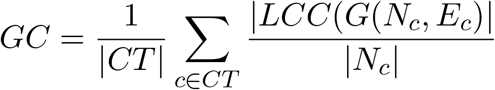

Where *CT* is the set of cell types, |*LCC*()| is the number of cells in the largest connected component. *G*(*N*_*c*_, *E*_*c*_) is the kNN subgraph for cell type *c* with *N*_*c*_ as the number of cells in cell type *c* and *E*_*c*_ denoting the edges in the kNN graph containing only the cells of type *c*.
- **k-nearest neighbor Batch Effect Test (***kBET* ): k-nearest neighbor batch effect test (*kBET*) (41) evaluates how well the label configuration of the k-nearest neighborhoods of cells match the global label configuration. *kBET* test is performed iteratively over a random subset of cells, and the value is calculated as the total rejection rates over all the iterations. As *kBET* works on the kNN graph, for the embedding-based output and corrected feature output methods, we have used *k* = 50 for calculating the kNN graph. *kBET* is applied to each batch separately to account for technical effects and changes in the cell type distribution across batches. For the kNN graphs containing disconnected components, *kBET* is calculated on each of the connected components. The number of nearest neighbors would differ for each cell in the graph-based method like BBKNN. So diffusion-based correction is used for graph-based output to get the same number of nearest neighbors for all the cells. *kBET* value is scaled between 0 to 1 so that the 0 value indicates poor batch mixing and 1 indicates perfect mixing of cells.

### Isolated label metrics

We adopted two isolated label scores from (23) to evaluate how well a method can capture rare cell identities. Isolated labels denote the cell types that are present in the least number of batches. For each of the isolated labels, isolated F1 and isolated ASW are calculated. These two metrics reveal how well the isolated cell types are separated from the rest of the cell types after integration.

For the calculation of the isolated F1 score, first we need to determine the clusters containing the particular isolated label. Lovain clustering is used for the clustering, and the resolution of the clustering algorithm is set such that the cluster has the largest number of isolated labels. The F1 score of isolated labels is then measured again the other cells in the cluster. F1 score ranges from 0 to 1 with 0 indicating that all the cells in the cluster are from cell types other than the isolated label and 1 indicating that all the cells are from isolated labels. Isolated label ASW is calculated considering the isolated cell type as one cluster and all other cell types in another cluster and separately computing their ASW.

### Other metrics

- **Graph inverse Simpson’s index LISI:** The inverse Simpson’s index is a diversity score that calculates the number of neighbors needed to be visited before appearing the same batch again. The value of this metric ranges from 1 to B (number of batches). This LISI score is used for the evaluation of both batch correction (iLISI) and bio conservation (cLISI) (15). To extend the LISI to the graph-based methods graph LISI (23) is used. In graph LISI, the distance over joint embeddings is replaced with the graph-based distance between the cells to get the large number of neighbors. Dijkstra’s shortest path algorithm is used to calculate the shortest distance between the cells. In cLISI, inverse Simpson’s index is calculated over the cell types i.e., the number of neighbors one needs to visit before getting the same cell type again in the neighborhood, whereas iLISI is calculated over the batches.
- **Proportion of positive and true positive cells:** We further adopted another single-cell level summary metric from (22) to evaluate the integration performance. For this, first, the cells are classified into positive and negative cells. Cells surrounded by the same cell type are classified as positive; otherwise, they are classified as negative. Considering the subset of positive cells only, a positive cell is classified as true positive if the distribution of batches around it is the same as the global batch distribution. The proportion of positive cells indicate the extent of biological conservation. In contrast, the true positive percentage indicates how well the batches are well mixed. These two proportions together serve as quantitative metrics for evaluating the batch effect removal performance.

### Composite scores

We have introduced two composite score metrics for a holistic comparison of the performance across all the metrics for biological conservation and batch correction respectively. Moreover, we computed a combined composite score that measures the average performance of a method in biological conservation and batch correction. Each composite score is computed by averaging the scaled values of all the metrics in that category. For a dataset, the scaled value for each individual metric is obtained through min-max normalization across all the competing methods.

The composite scores are calculated as follows:

1. 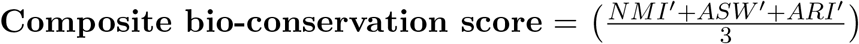
2. 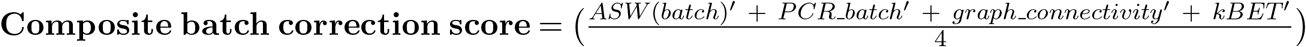
3. 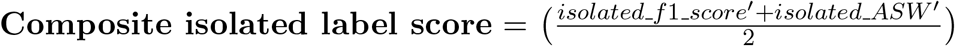
4. 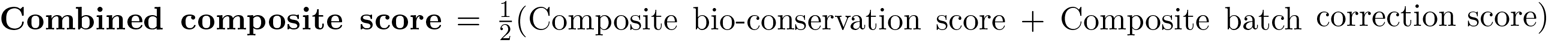

For any metric 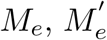 denotes the min-max scaled value across all the competing methods.

### Competing integration methods

We have compared our method against six state-of-the-art integration methods: scVI (20), Scanaroma (16), Harmony (15), Seurat (17), BBKNN (18), and INSCT (42). The details about the run configuration for these methods are provided in Supplementary Table 3.

## Data availability

The sources of the Human pancreas data are GSE81076, GSE85241, GSE86469, GSE84133, GSE81608 (Gene Expression Omnibus (GEO)) and E-MTAB-5061 (ArrayExpress). The dataset is also available at https://figshare.com/articles/dataset/Benchmarking_atlas-level_data_integration_in_single-cell_genomics_-_integration_task_datasets_Immune_and_pancreas_/12420968 (23). The source of Immune cell bone marrow are GSE120221 and GSE107727 (GEO). The immune cell peripheral blood 10X data were obtained from https://support.10xgenomics.com/single-cell-gene-expression/datasets/3.0.0/pbmc_10k_v3, GSE115189, GSE128066 and GSE94820. For the lung integration task, the Drop-seq data are available from GEO (GSE130148).

The processed Human Immune and Lung dataset are available at https://figshare.com/articles/dataset/Benchmarking_atlas-level_data_integration_in_single-cell_genomics_-_integration_task_datasets_Immune_and_pancreas_/12420968 (23).

The source of the Macaque Retina dataset is https://singlecell.broadinstitute.org/singlecell/study/SCP212/molecularspecification-of-retinal-cell-types-underlying-central-and-peripheral-vision-in-primates#study-download. The primary sources of Human and Mouse cell Atlas are https://figshare.com/articles/dataset/MCA_DGE_Data/5435866 and https://figshare.com/articles/dataset/HCL_DGE_Data/7235471. The processed Human-Mouse cell Atlas data used in our study is available at https://github.com/lkmklsmn/insct/tree/master/reproducibilty (42). All the experimental biological datasets used in this study are openly available and can be referenced from Supplementary Table 1.

## Code availability

The source code and usage tutorial for scDREAMER are freely available at https://github.com/Zafar-Lab/scDREAMER.

## Acknowledgements

This work was partially supported by the Science and Engineering Research Board (SERB), Government of India (SRG/2020/001333) and IIT Kanpur initiation grant (IITK /CS /2019236) to H.Z.

## Author contributions

H.Z. designed the study, A.S., M.K.P., and H.Z. developed the model and algorithm. A.S. and M.K.P. implemented the software and performed all experiments. All authors wrote and approved the manuscript.

## Competing interests

The authors declare no competing interests.

## Notes

### Competing Interest Statement

The authors have declared no competing interest.

