## Supplementary Information for "scDREAMER: atlas-level integration of single-cell datasets using deep generative model paired with adversarial classifier"

July 12, 2022

### Supplementary Figures

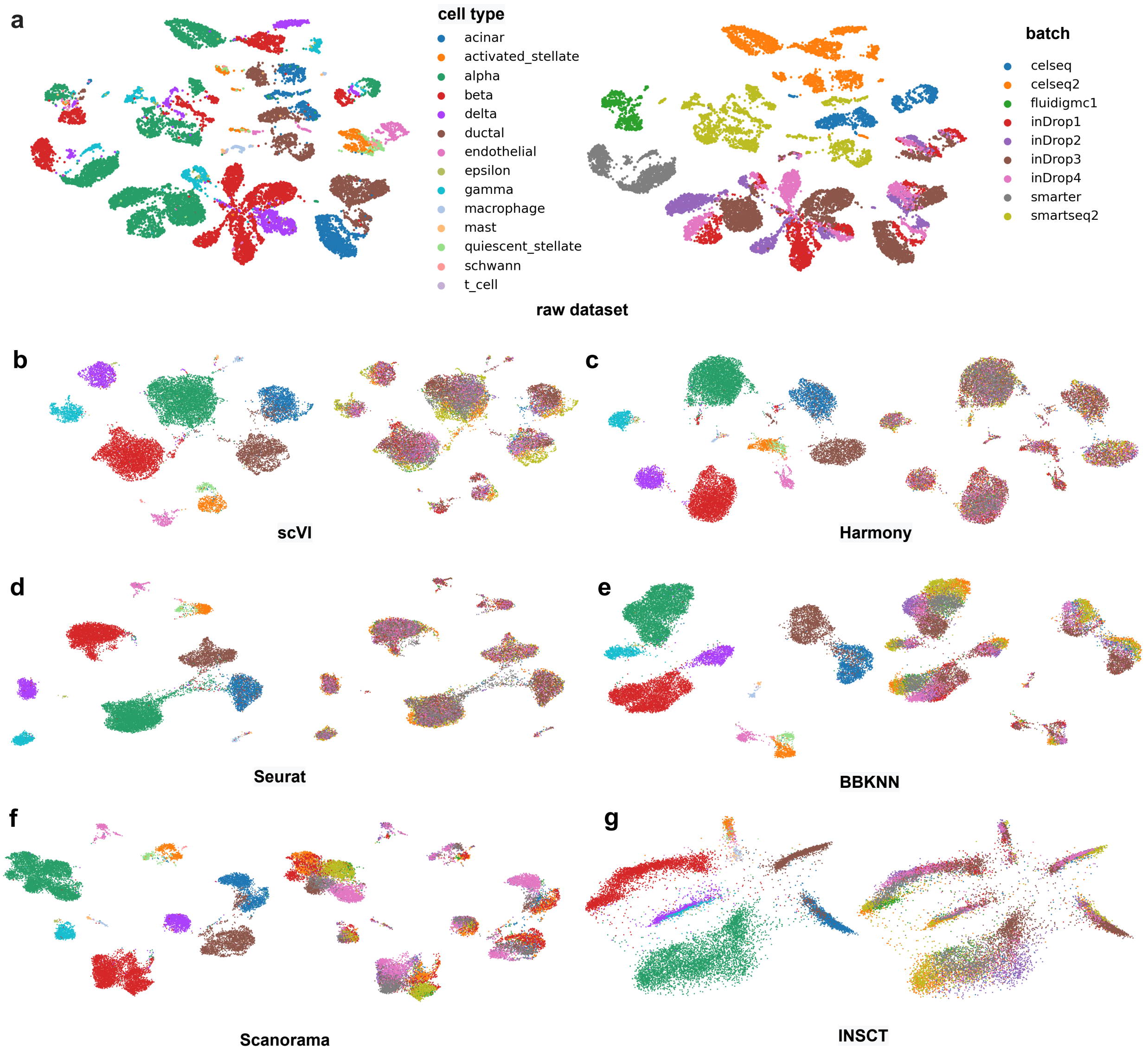

Supplementary Figure 1: Visualization of latent space embeddings for Human Pancreas Integration: a) Visualization of un-integrated human pancreas data annotated and coloured by different pancreatic cell types (left) and batch information (right). The dataset is generated from different single-cell sequencing protocols. Visualization of latent space embeddings post integration by different integration algorithms: b) scVI, c) Harmony, d) Seurat, e) BBKNN, f) Scanorama and g) INSCT.

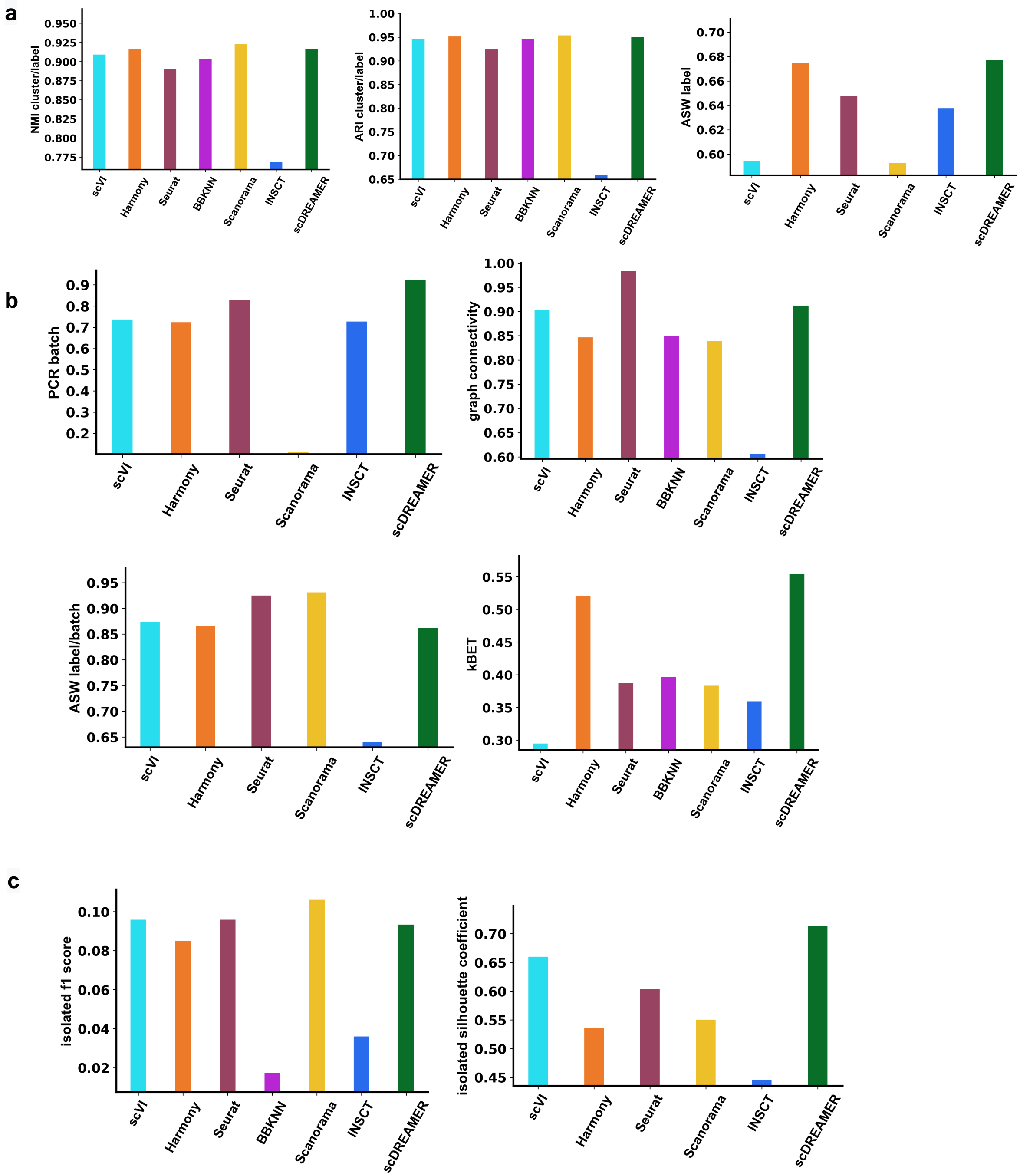

Supplementary Figure 2: Quantitative assessment of different methods for Human Pancreas Integration: a) Comparison of bio-conservation metrics i.e. NMI, ARI and ASW across different integration algorithms i.e. scVI, Harmony, Seurat, BBKNN, Scanorama, INSCT and scDREAMER. b) Comparison of batch-correction metrics i.e. PCR batch, graph connectivity, ASW label/batch and kBET across different integration algorithms. c) Comparison of isolated f1 score and isolated silhouette coefficient metrics across different integration algorithms.

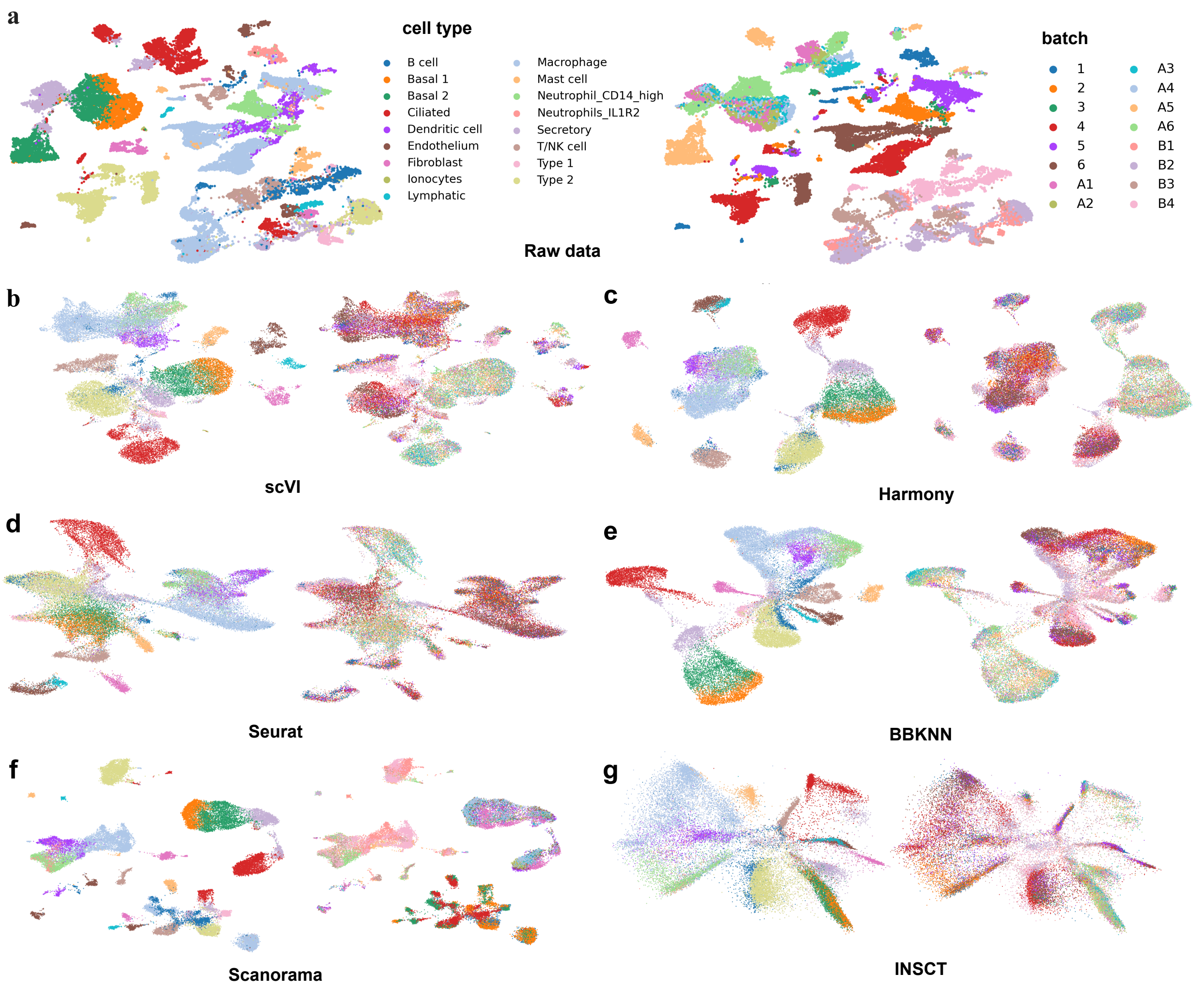

Supplementary Figure 3: Visualization of latent space embeddings for Lung Integration: a) Visualization of un-integrated Lung data annotated and coloured by different cell types (left) and batch information (right). Visualization of latent space embeddings post integration by different integration algorithms: b) scVI, c) Harmony, d) Seurat, e) BBKNN, f) Scanorama and g) INSCT.

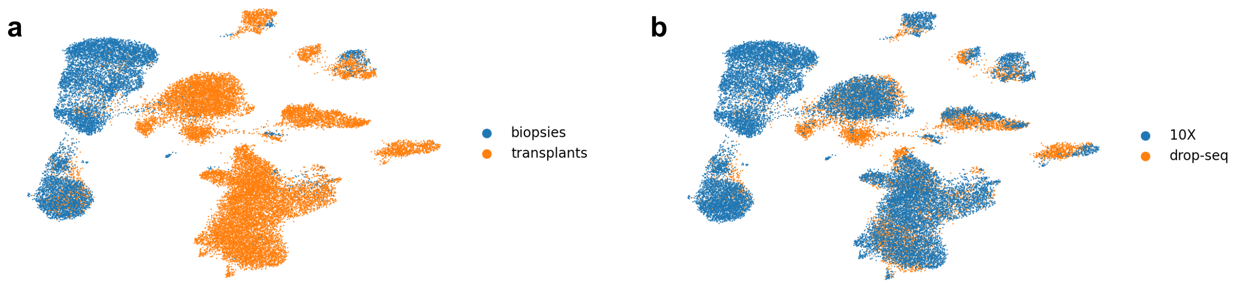

Supplementary Figure 4: Visualization of latent space embeddings inferred by scDREAMER for lung atlas integration with different annotations: a) Visualization of latent space embeddings annotated using different sampling locations. b) Visualization of latent embeddings annotated using different sequencing techniques.

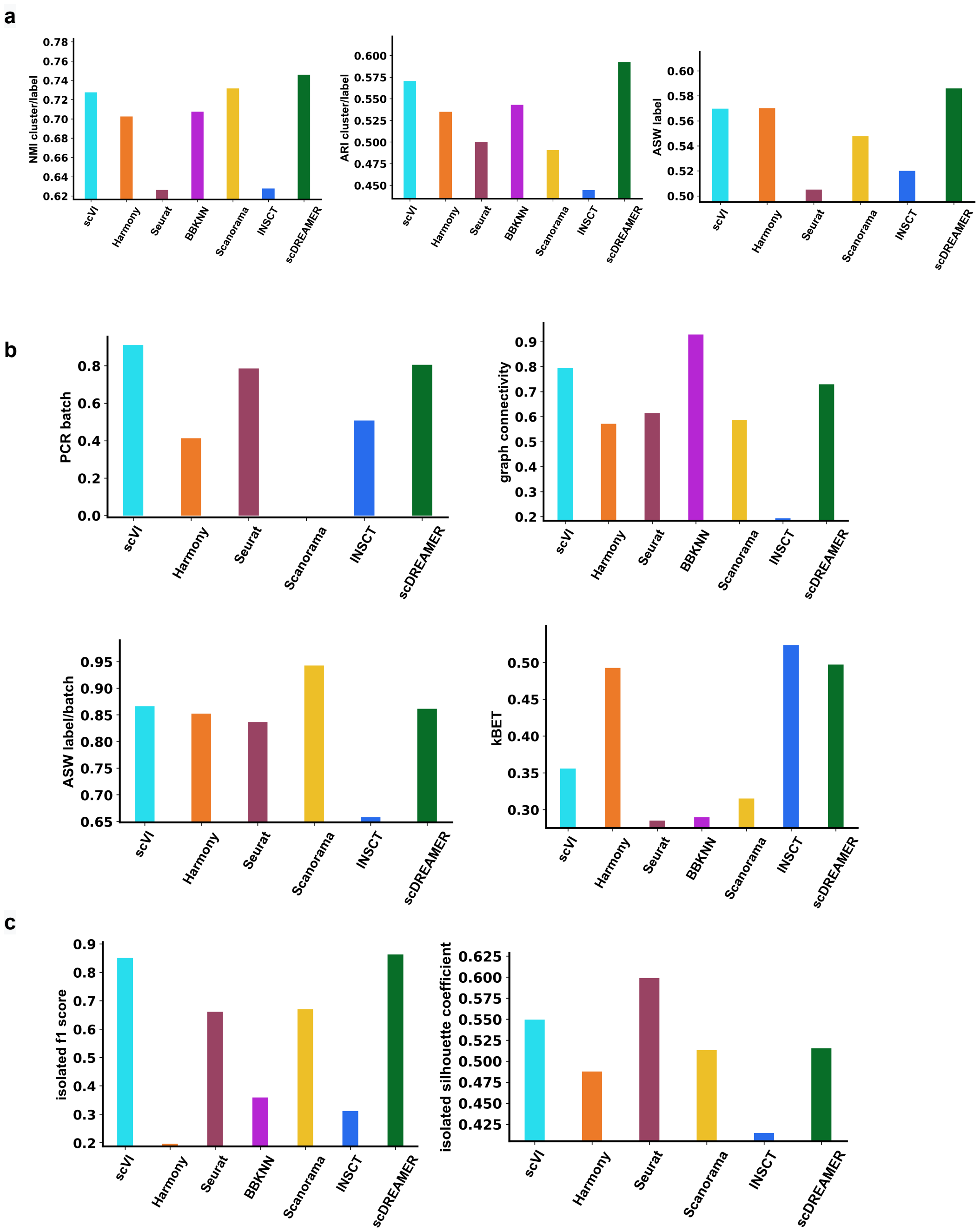

Supplementary Figure 5: Quantitative assessment of different methods for Lung Atlas Integration: a) Comparison of bio-conservation metrics i.e. NMI, ARI and ASW across different integration algorithms i.e. scVI, Harmony, Seurat, BBKNN, Scanorama, INSCT and scDREAMER. b) Comparison of batch-correction metrics i.e. PCR batch, graph connectivity, ASW label/batch and kBET across different integration algorithms. c) Comparison of isolated f1 score and isolated silhouette coefficient metrics across different integration algorithms.

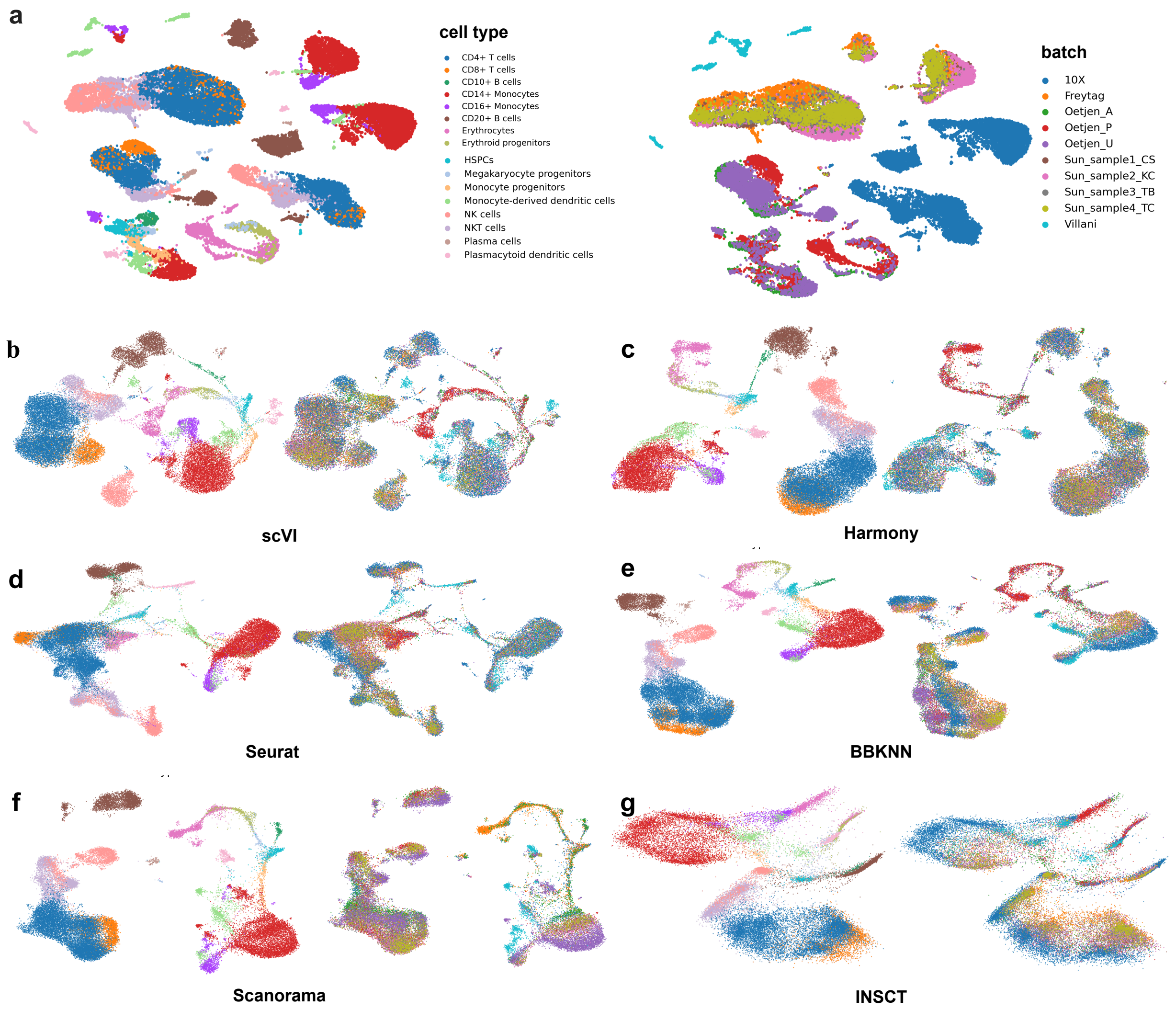

Supplementary Figure 6: Visualization of latent space embeddings for Human Immune Integration task: a) Visualization of un-integrated human immune dataset annotated and coloured by different immune cell types (left) and batch information (right). Visualization of latent space embeddings post integration by different integration algorithms: b) scVI, c) Harmony, d) Seurat, e) BBKNN, f) Scanorama and g) INSCT.

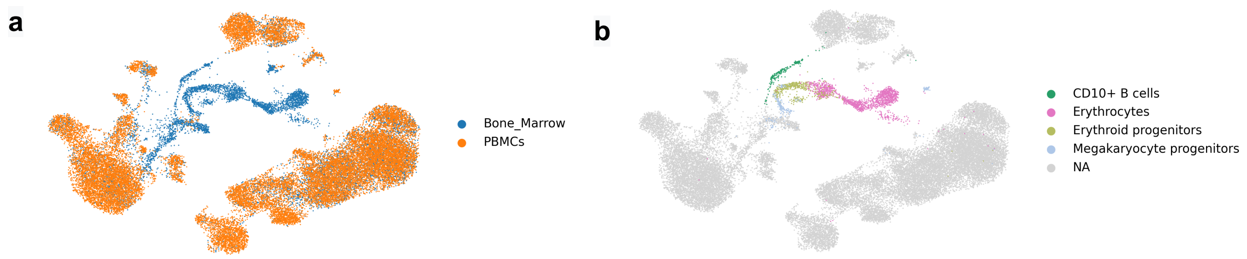

Supplementary Figure 7: Visualization of latent space embeddings inferred by scDREAMER for Human Immune Integration with different annotations: a) Visualization of Human Immune dataset annotated by tissue i.e. PBMC and Bone marrow b) Visualization highlighting the cell types in the trajectory (bone marrow specific) captured by scDREAMER.

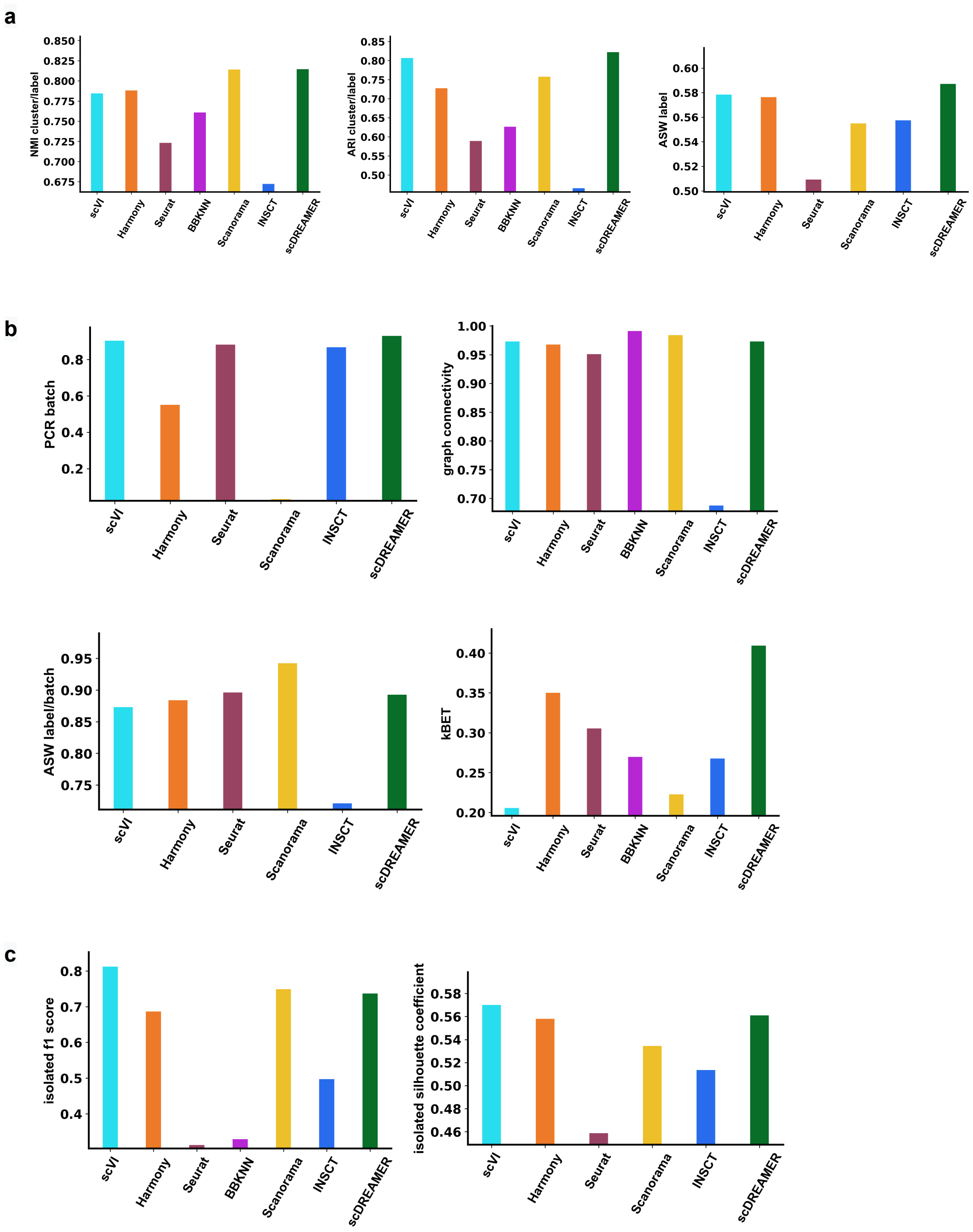

Supplementary Figure 8: Quantitative assessment of different methods for Human Immune Integration: a) Comparison of bio-conservation metrics i.e. NMI, ARI and ASW across different integration algorithms i.e. scVI, Harmony, Seurat, BBKNN, Scanorama, INSCT and scDREAMER. b) Comparison of batch-correction metrics i.e. PCR batch, graph connectivity, ASW label/batch and kBET across different integration algorithms. c) Comparison of isolated f1 score and isolated silhouette coefficient metrics across different integration algorithms.

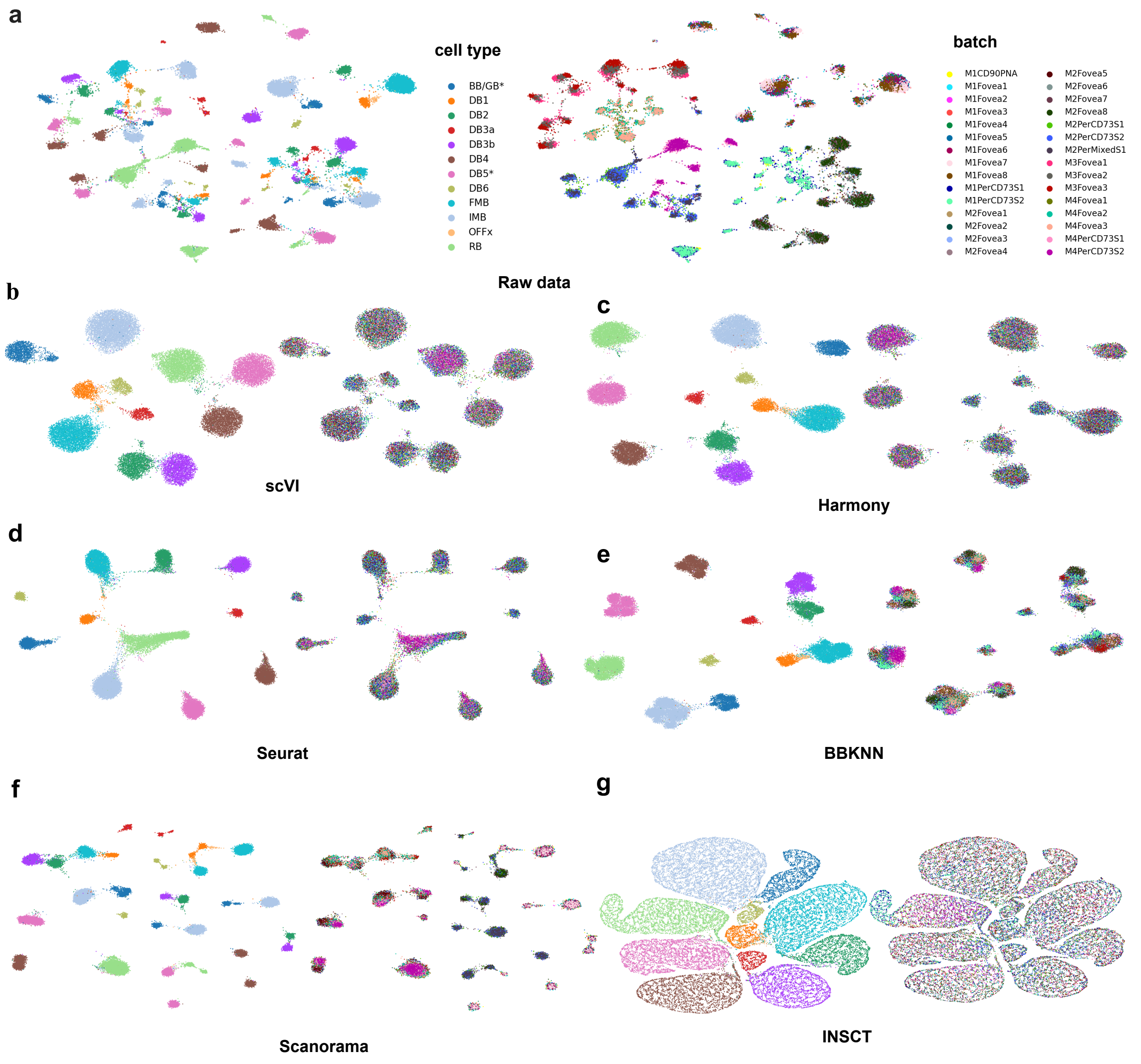

Supplementary Figure 9: Visualization of latent space embeddings for Macaque Retina Integration: a) Visualization of un-integrated macaque retina cells annotated and coloured by different cell types (left) and batch information (right). Visualization of latent space embeddings post integration by different integration algorithms: b) scVI, c) Harmony, d) Seurat, e) BBKNN, f) Scanorama and g) INSCT.

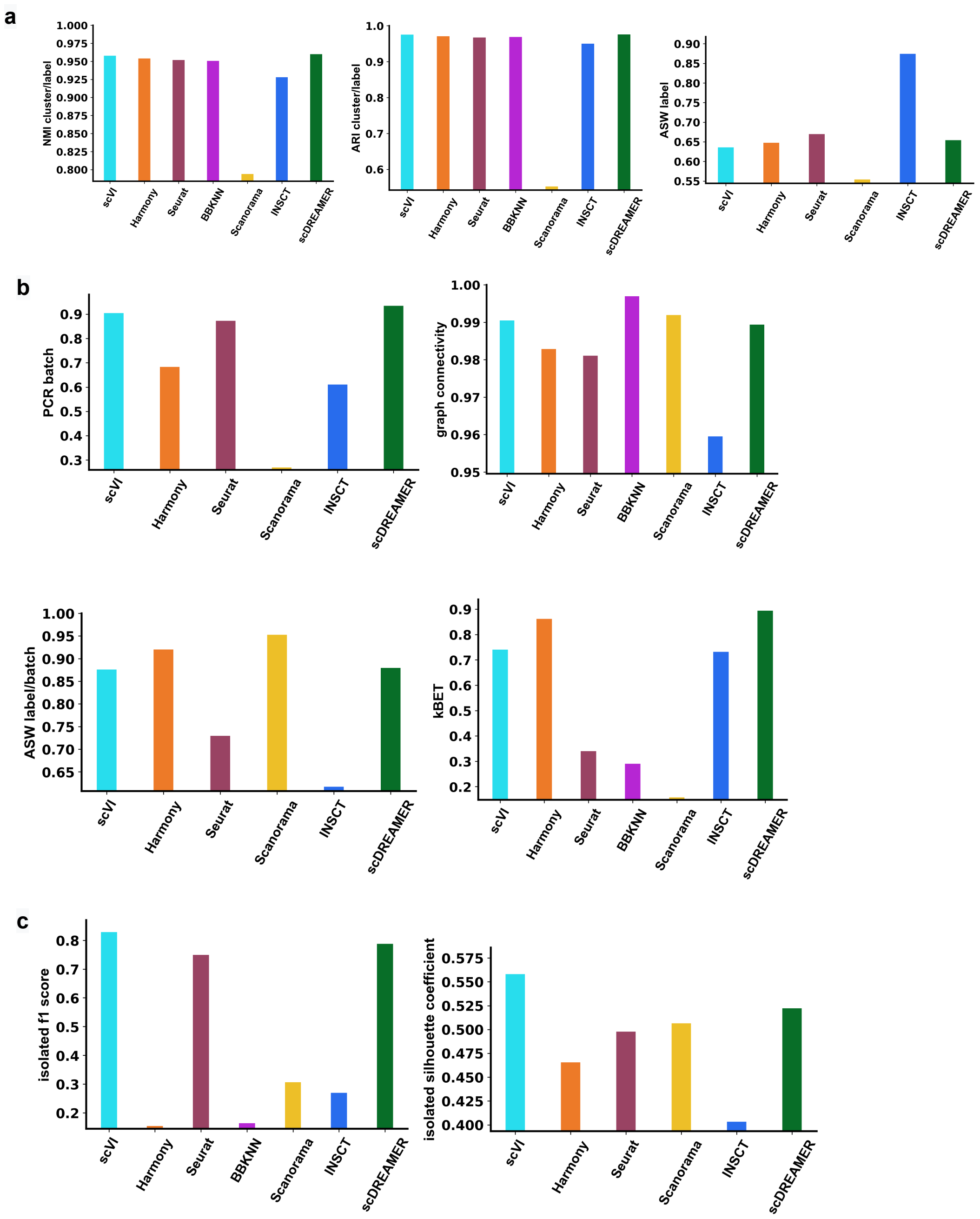

Supplementary Figure 10: Quantitative assessment of different methods for Macaque Retina Integration: a) Comparison of bio-conservation metrics i.e. NMI, ARI and ASW across different integration algorithms i.e. scVI, Harmony, Seurat, BBKNN, Scanorama, INSCT and scDREAMER. b) Comparison of batch-correction metrics i.e. PCR batch, graph connectivity, ASW label/batch and kBET across different integration algorithms. c) Comparison of isolated f1 score and isolated silhouette coefficient metrics across different integration algorithms.

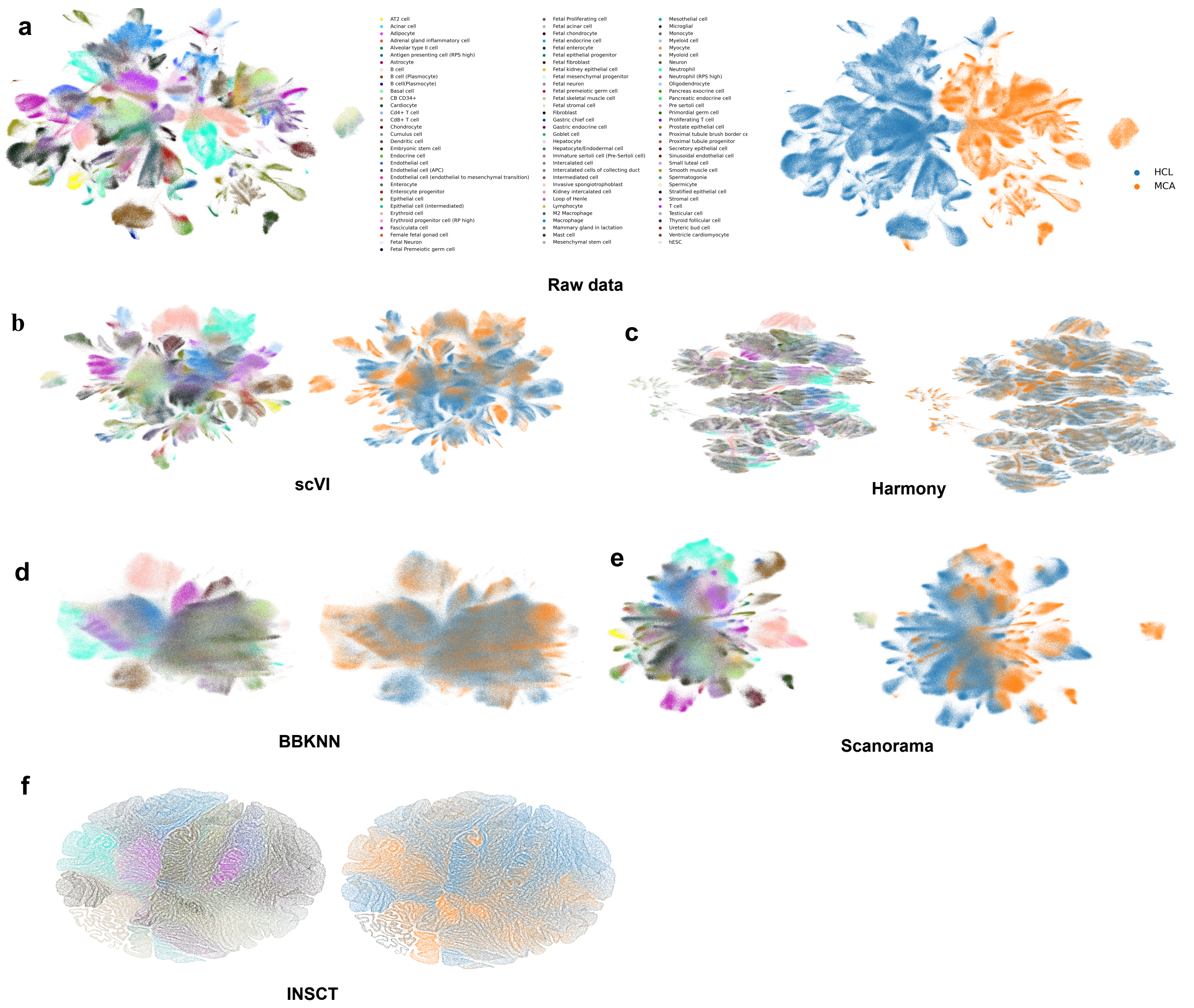

Supplementary Figure 11: Visualization of latent space embeddings for Human Mouse Integration: a) Visualization of un-integrated Human Mouse data annotated and coloured by different cell types (left) and batch information (right). The integration is performed across species i.e. human and mouse. Visualization of latent space embeddings post integration by different integration algorithms: b) scVI, c) Harmony, d) BBKNN, e) Scanorama, and f) INSCT.

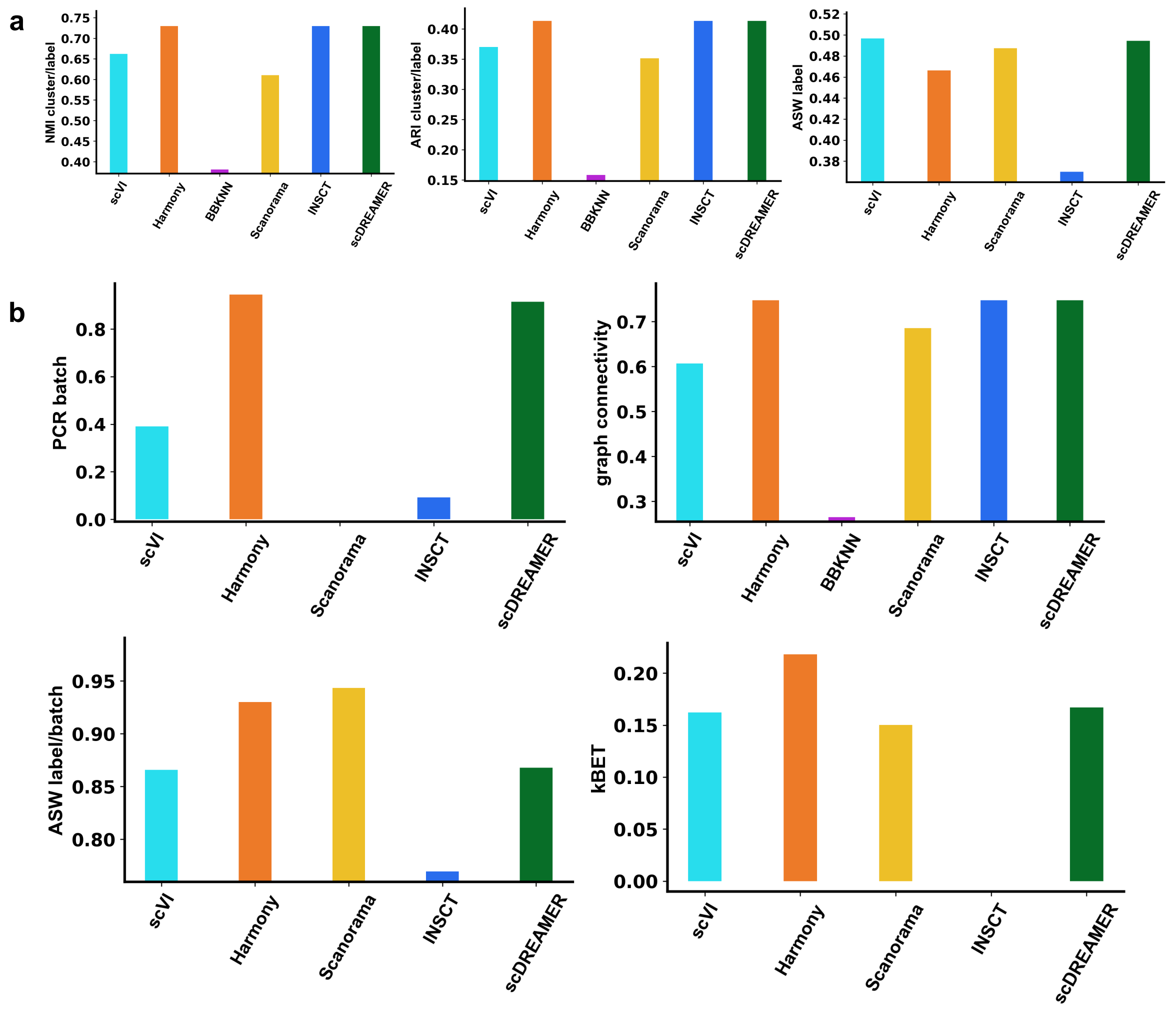

Supplementary Figure 12: Quantitative assessment of different methods for Human Mouse Integration: a) Comparison of bio-conservation metrics i.e. NMI, ARI and ASW across different integration algorithms i.e. scVI, Harmony, Seurat, BBKNN, Scanorama, INSCT and scDREAMER. b) Comparison of batch-correction metrics i.e. PCR batch, graph connectivity, ASW label/batch and kBET across different integration algorithms. c) Comparison of isolated f1 score and isolated silhouette coefficient metrics across different integration algorithms.

### Supplementary Tables

Supplementary Table 1: Description of the datasets used for benchmarking

| Dataset | Dimensions (cells * genes) | batches | cell types | Integration across | Source |
|---|---|---|---|---|---|
| Human Pancreas | 16382 * 34363 | 9 | 14 | Seq. protocols | link |
| Lung | 32472 * 15148 | 16 | 17 | Human donors; Laboratories | link |
| Immune Human | 33506 * 12303 | 10 | 16 | Tissue, seq. protocols | link |
| Macaque Retina | 30302 * 18323 | 30 | 12 | Macaques; Regions | link |
| Human Mouse | 933704 * 4999 | 2 | 97 | Species | Mouse, Human and link |

#### Pancreas

The Pancreas dataset comprises of data from 9 batches with dimensions 16382*2000 (cells *genes). Here, the batches denote different sequencing techniques i.e. celseq, celseq2, fluidigmc1, inDrop1, inDrop2, inDrop3, inDrop4, smarter and smartseq2. Also, there are 14 pancreatic cell types included in the dataset, namely, acinar, activated_stellate, alpha, beta, delta, ductal, endothelial, epsilon, gamma, macrophage, mast, quiescent_stellate, schwann and t_cell. The source of the dataset is https: //figshare.com/articles/dataset/Benchmarking_atlaslevel_data_integration_in_singlecell_genomics

#### Lung

The Lung dataset comprises of scRNA-seq data from 16 donor batches with dimension 32472 * 15148 (cells * genes). The dataset from all batches covers 17 different cell type including B cell, Basal 1, basal 2, ciliated, dendritic cell, endothelium, fibroblast, monocytes, lymphatic, macrophage, mast cell, secretory, T/NK cell, Type 1 cells, type 2 cells, neutrophil_CD144_high and neutrophils1L1R2 cells. The source of the dataset is https: //figshare.com/articles/dataset/Benchmarkingatlas−leveldataintegrationinsingle−cellgenomics

#### Immune Human

The Immune Human dataset comprises of data from 10 batches with dimensions 33506 * 12303 (cells * genes). The batches are 10X, Freytag, Oetjen_A, Oetjen_P, Oetjen_U, Sunsample1_CS,

Sunsample2KC, Sunsample3TB, Sunsample4_TC and Villani. Also, the dataset comprises of 16 different cell types i.e.CD4+ T cells, CD8+ T cells, CD10+ B cells, VD14+ monocytes, CD16+ monocytes, CD20+ B cells, Erythrocytes, Erythroid progenitors, HSPCs, Megakaryocyte progenitors, monocyte progenitors, monocytederived dendritic cells, NK cells, NKT cells, Plasma cells and plasmacytoid dendritic cells. The source of the dataset is https://figshare.com/articles/dataset/Benchmarking_atlaslevel_data_integration_in_singlecell_genomics

#### Macaque Retina

The Macaque Retina dataset comprises of data from 30 batches with dimensions 30302 × 18323 (cell * genes). The data from all batches covers 12 subclusters, namely, BB/GB*, DB1, DB2, DB3a, DB3b, DB4, DB5*, DB6, FMB, IMB, OFFx and RB. The batche samples include M1CD90PNA,M1Fovea(1-8), M1PerCD73S1, M1PerCD73S2, M2Fovea(1-8), M2PerCD73S1, M2PerCD73S2, M2PerMixedS1, M3Fovea(1-3), M4Fovea(1-3), M4PerCD73S1, M4PerCD73S2. The source of the dataset is https://singlecell.broadinstitute.org/single$_{c}ell/study/SCP212/molecular - specification - of - retinal - cell - types - underlying - central - and - peripheral - vision - in - primatesstudy - download$

#### Human Mouse

The Human Mouse dataset comprises of data from 2 batches i.e. human and mouse with dimensions 933704 × 4999 (cells * genes). There are total 92 cell types, majorly covering, B cell, Acinar cells, mesothelial cells neutrophil cells, endothelial cells, epithelial cells, thyroid cells and Endocrine cells.The source of the mouse atlas data is https://figshare.com/articles/dataset/MCA_DGE_Data/5435866 and the source of human atlas data is https: //figshare.com/articles/dataset/HCL_DGE_Data/7235471.

Supplementary Table 2: Parameter settings used for scDREAMER training

| **Hyper parameters** | **Hyper parameter values** |
|---|---|
| $\beta$, kl-scaling factor | 0.001 |
| $\eta$,learning rate | 0.0007 |
| $\eta_1$, learning rate (ELBO Loss) | 0.0002 |
| Optimizer used | ADAM optimizer |
| Batch size | 128, 256 (Human Mouse data) |
| $hvg$, highly variable genes of sc-RNA data | 2000 |
| Epochs | $300 - 400$ |
| $z_{dim}$, latent space dimensions | 10 |
| **Network** | **Architecture** |
| Encoder network | $hvg \longrightarrow 512 \longrightarrow 256 \longrightarrow z_{dim}$ |
| Decoder network | $z_{dim} \longrightarrow 256 \longrightarrow 512 \longrightarrow 3 \times hvg\ (x, \mu_x, \theta_x)$ |
| Batch-Classifier network | $z_{dim} \longrightarrow 256 \longrightarrow 512 \longrightarrow \lvert S \rvert$ (no. of batches) |
| Discriminator network | $hvg \longrightarrow 256 \longrightarrow 512 \longrightarrow 1$ |

### Other integration methods in detail

We have compared scDREAMER against six state-of-the-art methods i.e. scVI (1), Scanorama (2), Harmony (3), Seurat (4), BBKNN (5) and INSCT (6). More details on the methods and configuration are as follows.

Supplementary Table 3: Configuration of the competing methods

| **Methods** | **Principle** | **Configuration** | **Github (https://github.com/)** |
|---|---|---|---|
| scVI | Conditional variational autoencoder | version: 0.7.0a5 (default params.) | YosefLab/scvi-tools link |
| Harmony | PCA + clustering-based correction | version: 0.0.5 (default params.) | immunogenomics /harmony link |
| Seurat | CCA + Mutual nearest neighbors | version: 4.0.6 (default params.) | satijalab/seurat link |
| BBKNN | KNN graph integration | version: 1.5.1 (default params.) | Teichlab/bbknn link |
| Scanorama | SVD + Mutual nearest neighbors | version: 1.7.1 (default params.) | brianhie/scanorama link |
| INSCT | Batch-aware triplet neural network | version : 0.0.1 (default params.) | lkmklsmn/insct link |
